# Perivascular cells function as mechano-structural sensors of vascular capillaries

**DOI:** 10.1101/2024.02.16.580564

**Authors:** Cristiane M. Franca, Maria Elisa Lima Verde, Alice Correa Silva-Sousa, Amin Mansoorifar, Avathamsa Athirasala, Ramesh Subbiah, Anthony Tahayeri, Mauricio Sousa, May Anny Fraga, Rahul M. Visalakshan, Aaron Doe, Keith Beadle, McKenna Finley, Emilios Dimitriadis, Jennifer Bays, Marina Uroz, Kenneth M. Yamada, Christopher Chen, Luiz E. Bertassoni

**Author notes:** Luiz E. Bertassoni Knight Cancer Precision Biofabrication Hub OREGON HEALTH & SCIENCE UNIVERSITY – OHSU 2720 S Moody Ave, Portland OR 97201 USA **E-mail:**. **Author Contributions:** C.M.F., M.E.L.V., A.S., A.A., R.S., M.S., M.A.F., R.M.V., J.B. and M.U. performed experiments; C.M.F., M.E.L.V., A.M., A.T. engineered and optimized the microfluidic devices; C.M.F and A.D. analyzed nanostring data; K.B and M.F contributed with flow cytometry; E.D and K.M.Y. contributed with atomic force microscopy experiments; C.M.F., K.M.Y., C.C. and L.E.B wrote the paper; L.E.B. conceptualized the paper and supervised the experiments. **Competing Interest Statement:** Authors declare no conflict of interest.

## Abstract

A wide range of conditions, including chronic inflammatory diseases and cancer, are characterized by the fibrotic microarchitecture and increased stiffness of collagen type I extracellular matrix. These conditions are typically accompanied by altered vascular function, including vessel leakiness, abnormal capillary morphology and stability. The dynamic cell-matrix interactions that regulate vascular function in healthy tissues have been well documented. However, our understanding of how the gradual mechanical and structural alterations in collagen type I affect vascular homeostasis remains elusive, especially as a function of the interactions between endothelial and perivascular cell with the altered matrix. Here we hypothesized that perivascular cells might function as mechano-structural sensors of the microvasculature by mediating the interaction between endothelial cells and altered collagen type I. To test that, we utilized an organotypic model of perivascular cell-supported vascular capillaries in collagen scaffolds of controlled microarchitecture and mechanics. Our results demonstrate that capillaries cultured in soft reticular collagen exhibited consistent pericyte differentiation, endothelial cell-cell junctions, and barrier function. In contrast, capillaries embedded in stiff and bundled collagen fibrils to mimic a more fibrotic matrix induced abluminal migration of perivascular cells, increased leakage, and marked expression of vascular remodeling and inflammatory markers. These patterns, however, were only observed when endothelial capillaries were engineered with perivascular cells. Silencing of *NOTCH3,* a mediator of endothelial-perivascular cell communication, largely re-established normal vascular morphology and function. In summary, our findings point to a novel mechanism of perivascular regulation of vascular dysfunction in fibrotic tissues which may have important implications for anti-angiogenic and anti-fibrotic therapies in cancer, chronic inflammatory diseases and regenerative medicine.

**Significance Statement:** The fibrotic alterations in extracellular matrix structure and mechanics that are common to many chronic and inflammatory conditions are often associated with a decrease in vascular homeostasis. The mechanisms regulating these abnormalities remain poorly understood. Here, we demonstrate that perivascular cells play a critical role in sensing progressive microarchitectural and mechanical changes occurring in the ECM, drastically altering vascular capillary morphology and barrier function, and exacerbating the production of inflammatory and remodeling markers. These results point to a previously unknown mechano-structural sensory mechanisms mediated by perivascular cells in vascular capillaries that may help elucidate the progression of many profibrotic conditions, and point to possible new targets for antiangiogenic and antifibrotic therapies in cancer, chronic inflammatory conditions and regenerative medicine.

## Introduction

Maintenance of microvascular homeostasis is orchestrated through a dynamic interplay between endothelial and perivascular cells with the extracellular matrix (ECM) (1). Collagen type I is the most abundant protein in the human ECM (2, 3), and both its microarchitecture and mechanics are crucial for maintaining tissue function and driving disease progression (4, 5). For example, increased collagen stiffness and bundled architecture are hallmark features of ECM fibrosis (6, 7), and are associated with chronic inflammatory diseases, aging, and a cancer-promoting stroma (8, 9). Both endothelial and perivascular cells have been reported to sense and respond to the mechanics and architectural features of the ECM (10–13). However, the specific mechanisms regulating the response of vascular capillaries to the collagen structural and mechanical changes that are observed in these fibrotic conditions remain poorly understood.

Perivascular cells are known to stabilize the vasculature (3, 14), and play a significant role in regulating vessel permeability (15, 16). Mural cells have also been implicated in profibrotic events associated with acute tissue injury (17), inflammation (18) and cancer progression (19, 20). Perivascular cells occupy a strategic position between the ECM and the abluminal side of the endothelium. They share the endothelial basement membrane, and extend processes across multiple cells, thus forming a two-way interface bridging the endothelium with the surrounding ECM (21). Here, we hypothesize that perivascular cells create a structural mechanism for transmission of architectural changes and mechanical forces that sense and respond to the changes in collagen fibrils surrounding vascular capillaries, thus pointing to perivascular cells as key mechano-structural sensors in the microvasculature.

To test this hypothesis, we utilized an organotypic model of perivascular cell-supported vascular capillaries in collagen type I (15, 22) with distinct fibrillar microarchitectures that were achieved by varying the temperature of collagen fibrillogenesis (23), without changing the collagen concentration. This resulted in a spectrum of collagen type I substrates ranging from a low-stiffness reticular (finer, more netlike) network, akin to a healthy ECM, to a stiffer bundled fibrillar mesh, mimicking a fibrotic tissue. Our results demonstrate that vascular abnormalities in response to collagen changes are minimal when endothelial capillaries are cultured without perivascular cells. On the other hand, perivascular cells mediate the formation of increasingly abnormal and leaky capillaries, marked by high expression of inflammation and remodeling markers in stiff bundled collagen. Our findings, therefore, point to a critical and previously unknown role for perivascular cells in determining the response of the vasculature to a dysregulated collagen ECM of fibrotic tissues.

## Results

### Perivascular cells regulate vascular capillary morphology as a function of collagen stiffness and architecture

To investigate the regulatory role of perivascular cells in forming vascular capillaries within collagen of varying stiffness and microarchitectures, we employed a microfluidic device (22, 24) composed of a single channel surrounded by collagen type-I cast into a polydimethylsiloxane (PDMS) mold (**Figure 1 A**). To control collagen fibril stiffness and architecture, we adjusted the collagen gelation temperature, resulting in distinct fiber diameters and porosity, while keeping adhesion ligand density constant, as previously described (23). Specifically, we polymerized rat tail collagen hydrogels at 4, 16, 21, and 37°C, where higher temperatures produced a finer and more compact reticular collagen network, and lower temperatures yielded a more porous matrix with a bundled fibril microarchitecture (**Figures 1 B, S1**).

**Figure 1.**
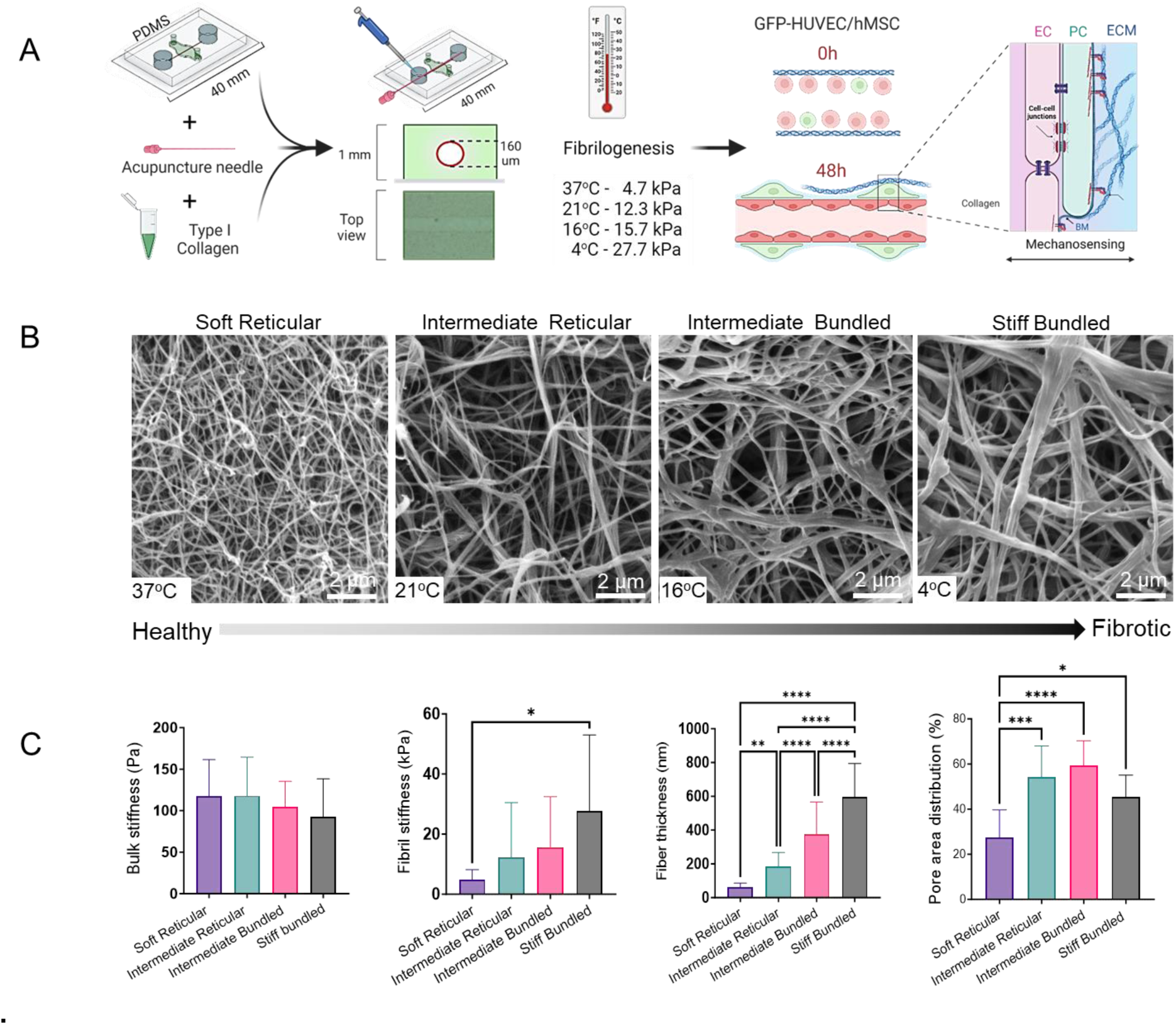
(A) Schematic diagram showing the steps to engineer perivascular-supported capillaries on-a-chip with different collagen stiffness and microarchitecture. (B) Representative scanning electron microscopy images of collagen mesh after crosslinking with different temperatures with gradual changes from soft reticular to stiff bundled. (C) Altering the crosslinking temperatures does not change the bulk modulus, but causes single collagen fibril elastic moduli to vary from ∼4.7 to ∼27.7 kPa, with progressive bundling of collagen fibers and heterogeneous pore distribution. \**P*<0.05, \*\**P*<0.01, \*\*\**P*<0.001, and \*\*\*\**P*<0.0001

Subsequently, we examined whether these structural differences correlated with fibril stiffness. When the hydrogel bulk modulus was measured via nanoindentation, no differences were found among groups (**Figure 1 C**), however, when we used conical-tipped pyramidal cantilevers with 1 µm tip size (23) to determine the stiffness of single-fibrils, differences in elastic modulus were significant (**Figures 1 B, S2**) and consistent with highly reticular fibers (37°C) having the lowest stiffness (4.7 kPa), and thicker fibrillar bundles (4°C) displaying significantly higher stiffness (27.7 kPa). Collagen crosslinked at 21°C and 16°C displayed intermediate stiffness of 12.3 kPa and 15.7 respectively, as illustrated in **Figure 1**. Moreover, as temperatures decreased, we observed an increase in fiber thickness and pore area distribution (**Figures 1 C**, **S1**).

We then examined the impact of collagen stiffness and microarchitecture on the formation of vascular capillaries by endothelial cells cultured alone in the engineered microchannels (**Figure 2**). We found that vessel formation occurred regardless of collagen stiffness, with minimal cell migration into the bulk collagen in all groups (**Figure 2 A, C**), with exception of a reduction in the number of sprouts from soft reticular to stiff bundled collagen (**Figures S3, S7**). Surprisingly, capillary morphology remained virtually unchanged, irrespective of collagen stiffness and architecture.

**Figure 2:**
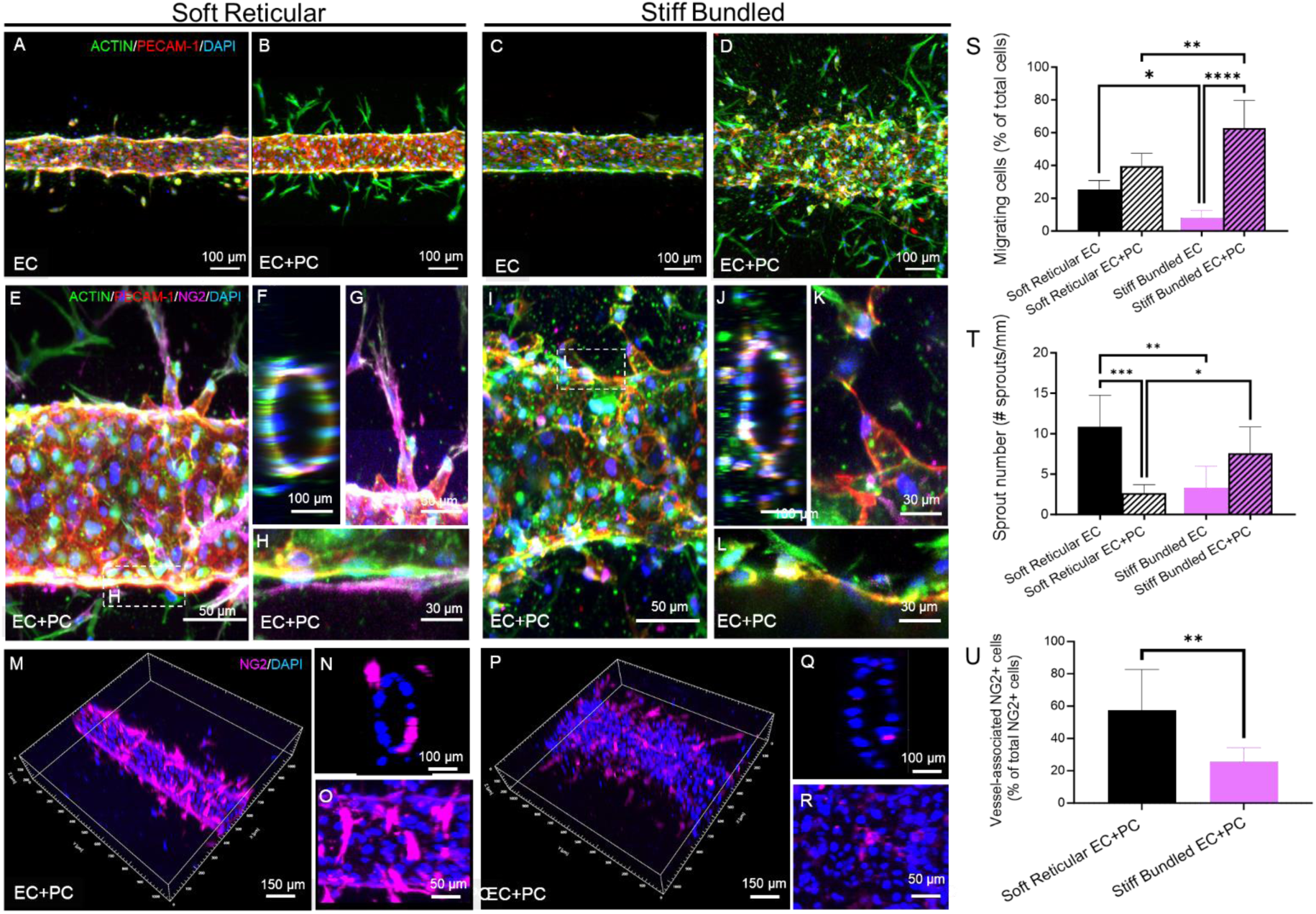
(A,C) Engineered capillaries with only endothelial cells (*EC*) presented similar morphologies despite collagen stiffness and microarchitecture. The presence of perivascular cells (*PC*) lead to more disrupted morphology and abluminal migrating cells in stiff bundled collagen (D,S). In addition, the presence of perivascular cells (B,D) drastically changed the morphology and number of capillaries within stiff bundled collagen leading to larger and more irregular vessels than in soft reticular collagen. (E-L, T) Capillaries engineered in both soft reticular and stiff bundled collagen showed angiogenic sprouts, however, only those in soft reticular fibrils had pericyte coverage (E-H,). (M-O) Capillaries engineered in soft reticular collagen also showed abundant pericyte coverage, while vasculature within stiff bundled collagen had fewer NG2+ cells associated with the endothelial wall (P-R, U). \**P*<0.05, \*\**P*<0.01, \*\*\**P*<0.001, and \*\*\*\**P*<0.0001

Next, we co-seeded HUVECs and hMSCs at a 4:1 ratio at a cell density of 8x10^6^ cells/mL in the microchannels. Within 24 hours, cells organized into an endothelialized capillary, consisting of an inner layer of endothelial cells surrounded by an outer layer of hMSCs (**Figure 2 E-L**), resembling the in-vivo morphology of pericyte cells in the vasculature (**Figure 2 M-O**). hMSCs within soft reticular collagen aligned longitudinally along the vessels and remained closely associated with the endothelial cells (**Figure 2 E-H, M-O, S**). Stiff bundled collagen led to increasing migration and proliferation of perivascular cells (**Figure 2 B,D**) towards the abluminal side of the vascular ECM (**Figure 2 I-L, P-R,S**). In addition, vessel morphology underwent significant changes as collagen became thicker in the stiff bundled group, with a significant increase in vascular diameter (**Figure S3**).

To assess whether collagen stiffness and microarchitecture influence the differentiation of hMSCs into a pericyte-like phenotype (2), we quantified NG2-positive cells adhered to the vessel wall. Our findings showed a consistent inverse correlation between stiffness and NG2 expression across all groups we tested (**Figures 2 M-R, S4, S5**). Accordingly, soft reticular collagen promoted pericyte differentiation near the vessel walls (**Figure 2 E-H,V**), while stiff bundled collagen had the opposite effect, showing the highest number of abluminal migrating cells and significantly fewer NG2+ cells (**Figure 2 I-L, P-R, V**). The presence of perivascular cells was also associated with a lower sprout length and number in soft reticular collagen, while stiff bundled fibrils had longer sprouts. This was opposite to the effects found in capillaries engineered with ECs only (**Figures 2 T, U, S6**).

### Perivascular cells regulate endothelial adhesion and barrier function in a collagen stiffness and architecture-dependent manner

To characterize endothelial cell adhesion as a function of stiffness and architecture, we again engineered capillaries with and without perivascular cells, and compared soft reticular collagen versus stiff bundled fibrils in vessels stained for PECAM-1/CD31 (Platelet and Endothelial Cell Adhesion Molecule) and N-cadherin (Neural-cadherin) (25). Interestingly, pericytes induced a significant increase in endothelial PECAM-1 expression within the soft reticular collagen, but not within the stiff bundled fibrils. In contrast, N-cadherin expression was prominently observed both in endothelial and perivascular cells in the stiff bundled group, while in the soft reticular group it was predominantly confined to perivascular cells. Notably, in the absence of perivascular cells, endothelial cells displayed limited N-cadherin expression (**Figure S8**) in fibrotic-like collagen.

To further understand the ECM-vasculature interactions in soft reticular and stiff bundled collagen, we characterized the expression of paxillin, a key component of focal adhesions that plays a role in the transduction of extracellular signals into intracellular responses, triggered by the engagement of integrins with the ECM. Upon integrin engagement with extracellular matrix, paxillin is phosphorylated, activating numerous signaling events associated with cell migration (26). While cells in the soft reticular collagen showed a sparser distribution of phosphorylated paxillin (p-pax), the stiff bundled collagen displayed enhanced p-pax that were concentrated adjacent to actin stress fibers **(Figure 3 E-F**). Instead, laminin, a critical basement membrane component, was significantly reduced only in EC and perivascular containing capillaries engineered within stiff bundled collagen (**Figure 3 G, J**).

**Figure 3:**
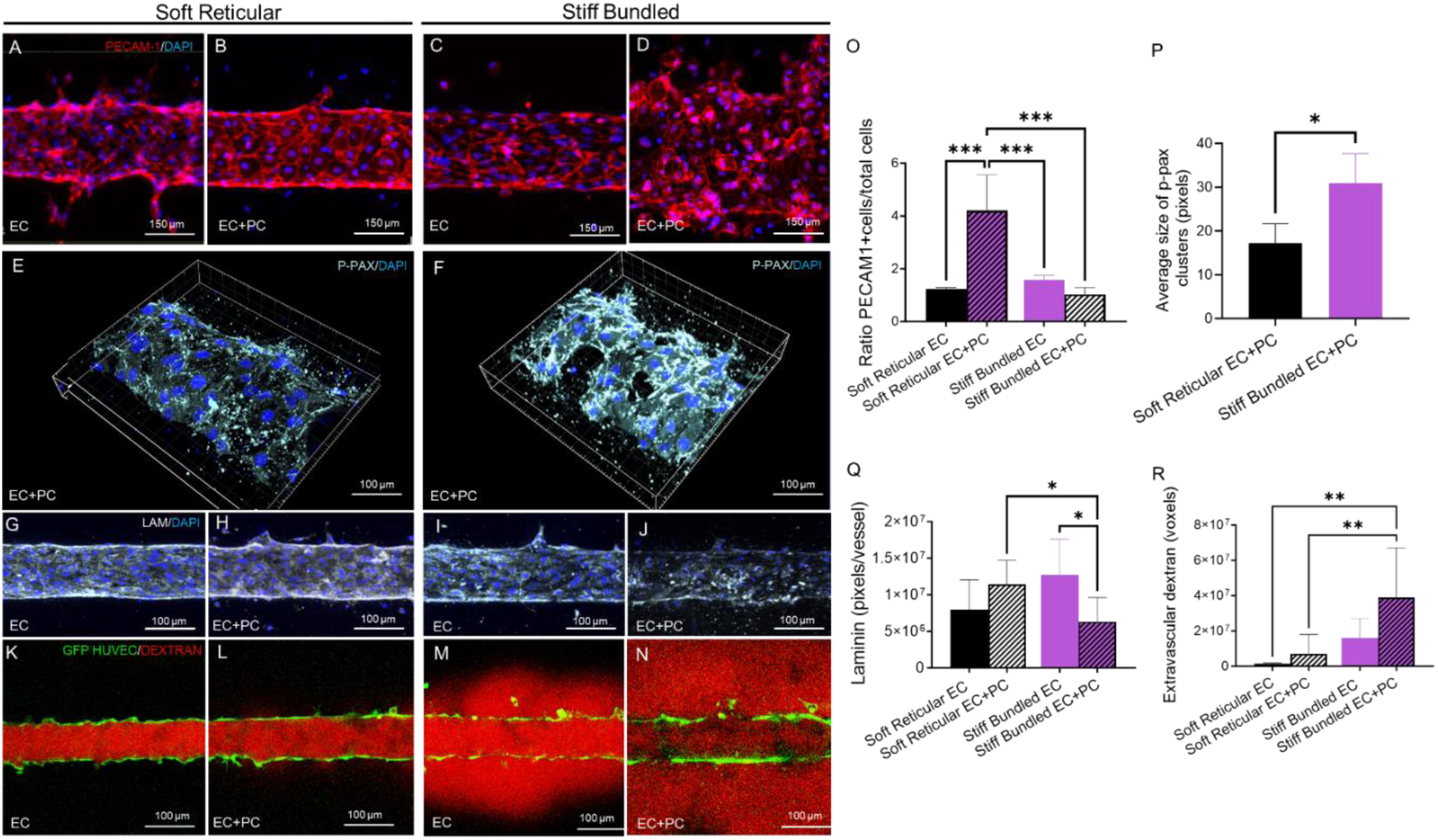
(A-D) Capillaries with only endothelial cells expressed PECAM-1 at comparable levels, while PECAM-1 is increased in capillaries with perivascular cells in soft reticular collagen. Heterogeneous and decreased expression of PECAM-1 is observed in capillaries engineered with perivascular cells in stiff bundled collagen. (E,F) Phosphorylated paxillin is overexpressed in vasculature engineered in stiff collagen. (G-J) Laminin was expressed in both groups regardless the absence of perivascular cells, however, perivascular cells in stiff bundled collagen showed irregular deposition of laminin across the capillary. (K-N) Vascular capillaries engineered in soft reticular collagen preserved barrier function regardless the presence of perivascular cells, while stiff bundled collagen was associated with reduced barrier function even in the presence of perivascular cells. \**P*<0.05, \*\**P*<0.01, \*\*\**P*<0.001, and \*\*\*\**P*<0.0001

Based on our observations that stiff bundled collagen drives perivascular-induced changes in endothelial adhesion, we examined if there was a change in barrier. Overall, variations in the anchorage and cell-cell adhesion systems led to sustained barrier function in vascular capillaries engineered with perivascular cells in soft reticular collagen, but disrupted barrier function in capillaries engineered with perivascular cells in stiff bundled collagen (**Figure 3 K-N**). Interestingly, the presence of perivascular cells maintained the barrier function even when collagen had intermediate stiffness of ∼12.3 kPa that was almost three times the stiffness of soft reticular collagen (∼4.7 kPa). However, such barrier function was not preserved when stiffness was ∼15.7 kPa, and the collagen mesh was arranged in bundles (**Figure S9**). The positive influence of a soft reticular ECM in endothelial cell-cell adhesion was only significant in the presence of perivascular cells. Similarly, the negative impact of a stiff bundled architecture on cell-matrix interactions and barrier function was only observed when perivascular cells were co-cultured with ECs. Together, these data point to the critical role of perivascular cells in mediating both structural and mechanical events in vascular capillaries.

### Perivascular cells in stiff bundled collagen modulate genes related to inflammation and remodeling

To further understand the role of perivascular cells in the morphological and functional differences observed between soft reticular and stiff bundled collagen, we screened 770 genes using a nanostring sequencing (**Dataset S1** and **Figure S10**). From the top 10 most differentially expressed genes in the vasculature engineered in stiff bundled collagen compared to that engineered in soft reticular collagen, we identified genes associated to inflammatory signaling (*CXCL8*), ECM synthesis and remodeling (*TP53, TIMP1, COL3A1, MMP1, DCN*), cell proliferation and differentiation (*TGFB1, ACVR1C, SFRP1*), and cell-cell junction (*CLDN*) (**Figure 4 A-C**). Enrichment scores showed that capillaries engineered in stiff bundled collagen activated pathways related to cell migration, endothelial, proliferation, lysyl oxidase remodeling, chemotaxis and cancer (**Figures 4 D, E, S10-14**). On the other hand, capillaries in soft reticular fibrils activated pathways related to cell junction organization, TGFB1 signaling and control of chemotaxis, cell adhesion, among other signaling (**Figures 4 D, E, S10-14**).

**Figure 4:**
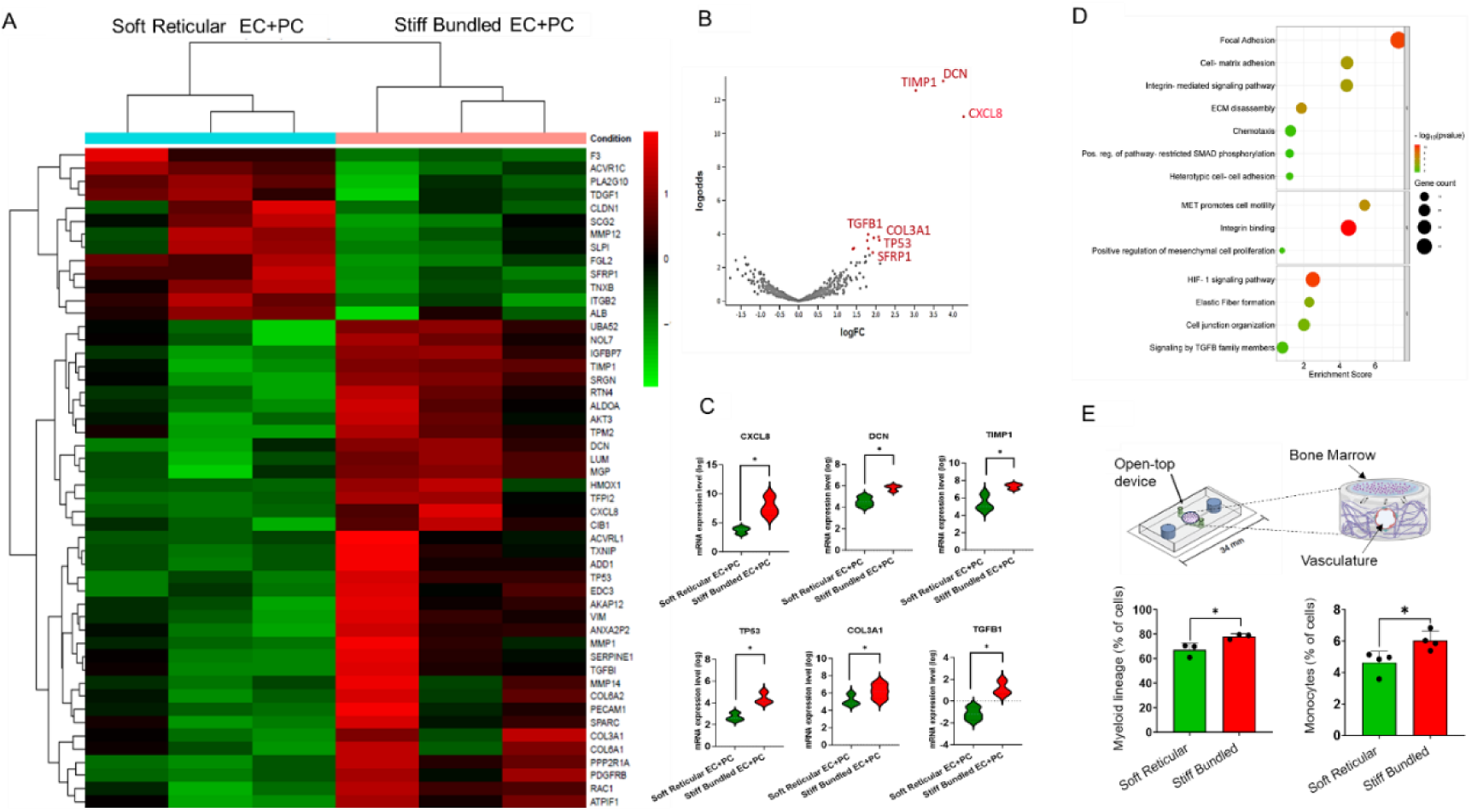
(A) Heat map comparing the effect of collagen stiffness and architecture on gene expression in the presence of perivascular cells. (B) Volcano plot showing that genes encoding for decorin, CXCL8 and TIMP1 were the most highly expressed by the vasculature in a stiff environment. (C) Pairwise comparison of the 9 most different log values for gene expression. (D) Enrichment score and pathways associated with capillaries engineered either in soft reticular or stiff bundled collagen. (E) Engineered bone marrow cells were seeded on top of the soft and stiff vasculature and allowed to interact for 3 days. In the presence of vasculature in stiff matrix, bone marrow cells tended to express more monocytic and myeloid markers consistent with CXCL8

To dissect the specific role of perivascular cells in a stiff bundled collagen matrix, we compared the expression data between capillaries with and without perivascular cells (**Figure S15**). We found that perivascular cells were particularly relevant in modulating genes related to the extracellular matrix such as *COL1A2, COL1A1, COL6A2, FN1, LOX, DCN, TIMP1*, and especially *CXCL8*. The presence of perivascular cells in a soft reticular environment led to the increase of *TGFB1* gene expression and pathways related to cell adhesion, integrin binding and ECM receptor interaction (**Figure S16**).

*CXCL8* was the most differentially expressed gene in the stiff bundled collagen when ECs were co-cultured with perivascular cells. Building upon the established connection of interleukin 8 (coded by the *CXCL8* gene) and inflammatory cell recruitment, we sought to further determine the effect of a stiff bundled microenvironment in vascular inflammatory paracrine signaling and chemotaxis from bone marrow cells. To that end, we prepared vascular channels using the same method utilized throughout the study, but in in open-top microdevices, and seeded an organotypic bone marrow model on top of the engineered capillaries (**Figure 4 E**). Cells in the engineered bone marrow had significantly higher differentiation into monocytic and myelocytic lineages in the presence of capillaries engineered in stiff bundled collagen, and similar differentiation into neutrophils, and hMSC (**Figures S17**, **S18**).

### Silencing Notch3 restores pericyte coverage, capillary morphology and barrier function

Our results showed marked differences between soft reticular and stiff bundled groups when vessels were engineered with perivascular cells. Moreover, our enrichment score analyses showed a significant effect of matrix adhesions and cell-matrix interactions (Figure 4D). Therefore, we first targeted the interaction between perivascular cells with the matrix by silencing the integrin B1 gene (*ITGB1*) in hMSCs before seeding the co-culture into the collagen channels. We hypothesized that blocking integrins would normalize the vasculature in the stiff bundled collagen. However, contrary to our expectation, not only vessels engineered in stiff bundled collagen did not normalize, but also the soft reticular collagen showed more migratory cells and less pericyte coverage, thus pointing to the importance of establishing adequate perivascular cell-matrix interaction irrespective of the matrix characteristics (**Figures S19, S20**). In other words, no perivascular cell adhesion to the matrix is worse than compromised perivascular cell adhesion to a stiff matrix. These results, therefore, suggested that our findings may be mediated at the cell-cell interface between perivascular and endothelial cells. To test that, we targeted NOTCH3, which was demonstrated to be a key mediator of the interaction between endothelial and perivascular cells (27). Our results showed that silencing *NOTCH3* decreased perivascular cell migration significantly in stiff bundled capillaries, bringing it to levels that were statistically comparable to soft reticular collagen (**Figure 5 A, E, I**), resulting in increased perivascular cell coverage. A similar effect was observed for NG2 expression (**Figure 5 C, G, K)**. In addition, *NOTCH3* silencing in perivascular cell significantly improved vessel barrier function in the stiff bundled group to levels that were comparable to those observed within soft reticular collagen (**Figure 5 A, E, J, insets**). Since IL-8 (*CXCL8* gene) was the most highly-expressed chemokine in the stiff bundled collagen group and silencing *NOTCH3* largely reversed the negative effects found in this group, we measured IL-8 levels in the supernatant of controls and silenced samples (**Figure 5 L**). Our results confirmed that the reversal in vascular morphology and function observed upon *NOTCH3* silencing was reflected in IL-8 paracrine signaling as well (**Figure 5 L**). Lastly, since *TGFB1* gene was overexpressed in the presence of perivascular cells both soft and stiff collagen groups, we also silenced *TGFB1* to probe its regulatory function in perivascular structural and mechanosensing. Notably, silencing *TGFB1* also normalized pericyte coverage and the number of migrating cells in the fibrotic vasculature (**Figures S19, S20**), although not as markedly as for *NOTCH3*. In addition, silencing *TGFB1* affected the capillaries engineered in the soft reticular group as well.

**Figure 5:**
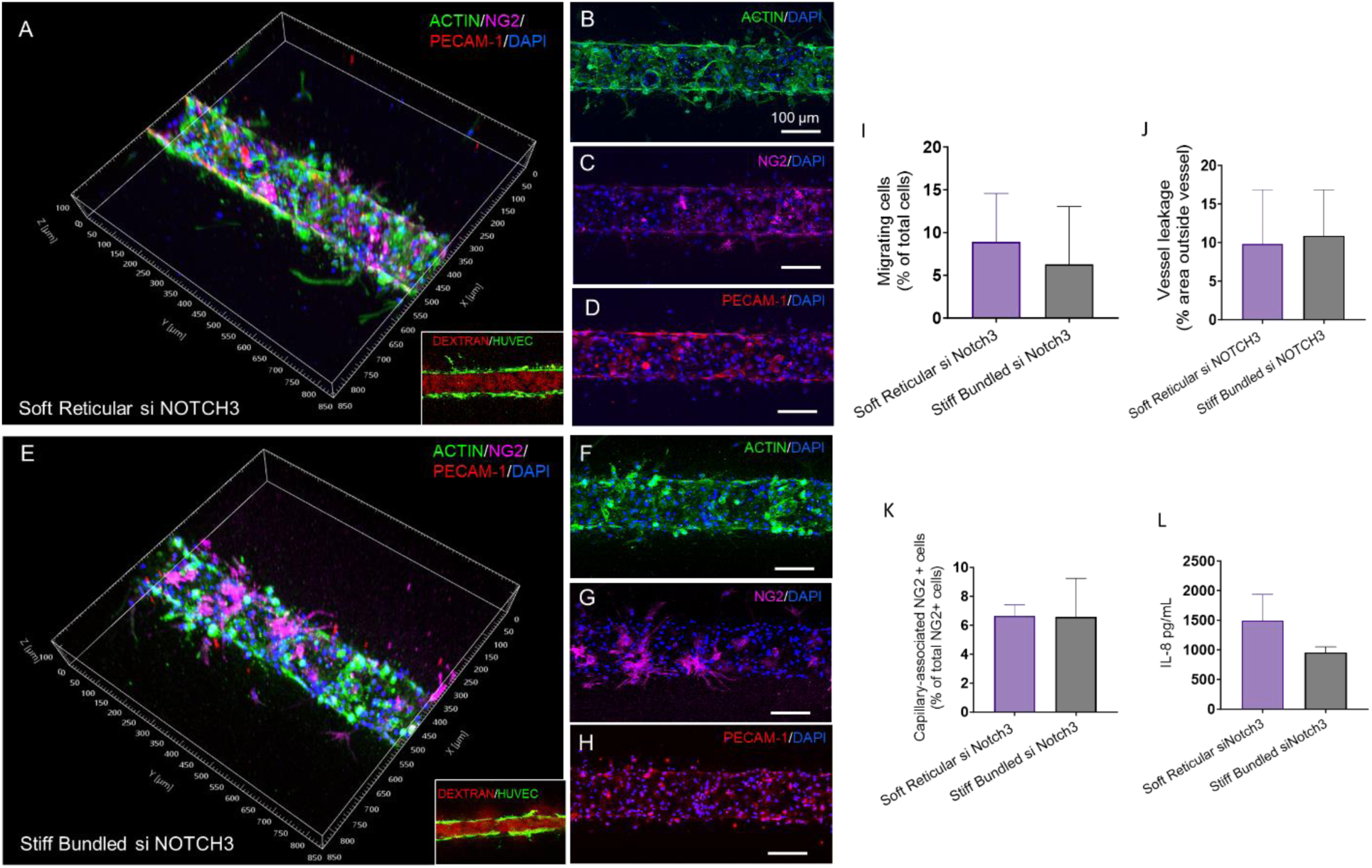
Normalization after Notch3 knockdown. Morphology (A,B, E,F), pericyte coverage (C,G) and PECAM-1 expression (D,H) were similar between vasculature engineered in a soft reticular (healthy) or stiff bundled (fibrotic) environment. (I,J,K) Migrating cells, pericyte coverage and vessel leakage was around 10%, similar in both groups. (L) In addition, interleukin 8 secretion was comparable for both groups. Non-silenced controls are in Figure S20.

## Discussion

Changes in the structure and mechanics of collagen are key hallmarks of fibrotic conditions, including many cancers, chronic and inflammatory diseases. However, the response of vascular capillaries to these dynamic alterations in the collagen ECM remain poorly understood. Here we demonstrate that perivascular cells are key mediators of the ability of capillaries to respond to collagen architectural and mechanical changes associated with fibrotic alterations around the vasculature. In the absence of perivascular cells, vascular morphology was not significantly altered, irrespective of changes in fibril thickness, stiffness and porosity **(Figure 2 A, C)**. Capillaries with perivascular cells, on the other hand, responded to a progressively more fibrotic collagen matrix with abnormal abluminal migration (**Figure 2 B, D**), lowered expression of endothelial adhesion markers (PECAM-1) (**Figure 3 A-D**) and pericyte differentiation (NG2) (**Figure 2 M-R**), becoming leakier **(Figure 3 K-N)** and more abnormally anchored to the matrix (**Figure 3 E-J**). Moreover, perivascular cells mediated the upregulation of inflammatory cytokines and matrix remodeling proteins in stiffer and more fibrotic collagen (**Figure 4 B, C**), dependent upon their communication with ECs (**Figure 5 A, E**). These results point to a possible feedback loop, where perivascular cells mediate vascular inflammation and remodeling, thus further exacerbating the ECM changes associated with the ongoing fibrotic alterations in the matrix.

Fibrotic tissues display a stiffer ECM, which is recognized to result from the thicker arrangement and density of individual collagen fibrils (6). We found that the expected pattern of endothelial monolayer formation, surrounded by tightly adhered pericytes was gradually disrupted as fibrils became stiffer, which is likely to result from the bundling of multiple fibrils and consequent increase in fibril thickness (**Figures S3, S4**) (8, 23). Two events are likely to be associated with this phenomenon. Firstly, stiffer matrices have been shown to reduce the differentiation of hMSCs into pericytes in co-cultures with ECs (11, 28). Secondly, poorly differentiated pericytes preserved the more migratory phenotype of hMSCs, and were more likely to penetrate the pore spaces in the collagen matrix, thus reducing the confinement of ECs to the engineered channels. Interestingly, that phenomenon was only present when perivascular cells were used, since EC-only capillaries did not show the same pattern of abluminal cell migration, despite the same characteristics of the collagen matrix. Of note, we also found correlation between the local expression of pericyte-associated markers and their distance from ECs for individual cells in the vessels, which suggests potential juxtacrine communication between ECs and perivascular cells mediating pericyte differentiation of hMSCs (**Figure S21**).

Since ECs and differentiated pericytes share the vascular basement membrane, we sought to determine if capillaries engineered in soft reticular collagen had a different pattern of cell-cell and cell-matrix adhesion than capillaries in stiff bundled fibrils. Paxillin, a key component of integrin signaling, requires tyrosine phosphorylation for integrin-mediated cytoskeletal reorganization and is a marker of activated focal adhesion kinases (26). The significantly more clustered pattern of phosphorylated-paxillin and overall lower expression of laminin in stiff bundled collagen, points to important alterations off the anchorage system displayed in a healthy-like ECM versus that of a fibrotic-like matrix (**Figure 3 E, F**). Endothelial cell-cell adhesion is mediated by vascular endothelial cadherin (VE-cad), while endothelial to perivascular cell contact is mediated by N-cadherin (N-cad) (29). N-cadherin is induced in perivascular cells in response to vascular injury in models of stiffening and proliferation (30), and both collectively mediate vascular barrier function (27, 30, 31). Similar to the pattern of expression of PECAM-1, phosphorylated-paxillin immunofluorescence revealed regions without significant expression (**Figure 3E,F**), indicating possible gaps in the cell-cell and cell-matrix communication system contributing to significantly leakier vessels in more fibrotic collagen. Likewise, the expression of N-cad followed a similar pattern of alterations in stiff bundled versus soft reticular collagen hydrogels.

Assessment of the 770 genes (**Figure S10 and Dataset S1**) expressed in capillaries with and without perivascular cells showed a marked increase in *CXCL8* expression, which encodes for interleukin 8 (IL-8), a canonical marker of inflammation, in both soft reticular and stiff bundled collagen. We hypothesized that high expression of *CXCL8* should lead to increased levels of IL-8, and consequently, an inflammatory response mediated by immune cells. Covering capillaries engineered in stiff bundled collagen with an organotypic model of bone marrow resulted in a significant increase in differentiation of human stem cells into monocytes and myeloid lineage cells (**Figure 4 E**). This supports the conjecture that IL-8, among other cytokines produced by the capillaries in a fibrotic matrix, may exacerbate the action of inflammatory cells in the perivascular microenvironment, such as monocytes, neutrophils and dendritic cells, which derive from a myeloid lineage (32), a finding that may have important implications for cancer progression and metastasis.

A proinflammatory microenvironment has been well established to accelerate matrix remodeling (33). Consistent with that notion, we also found that decorin (*DCN*), a matrix proteoglycan that is important for collagen fibril assembly (34), *COL3A*, which encodes new collagen production, and tissue inhibitor of matrix metalloproteinase (*TIMP*), which contributes to matrix degradation, were both significantly upregulated with *TP53*, which is known as the master regulator of vascular remodeling (**Figure 4 B,C**) in fibrotic collagen (35, 36). These findings indicate that a fibrotic matrix may induce inflammation, thereby unbalancing the expected process of vascular remodeling that guarantees the natural homeostasis of healthy vasculature. Our data suggest that this may occur by promoting abnormal secretion of new matrix and preventing controlled degradation of abnormal matrix proteins. Interestingly, these conditions fit within the well-reported progression of fibrotic diseases, where an increase in matrix stiffness is accompanied by inflammation and a worse prognosis or clinical outcomes, such as in breast cancer (37), diabetes (38, 39), and cardiovascular diseases (40, 41). More importantly, this points to a potential target for cancer therapy (42), where the gradual fibrosis of the premalignant microenvironment that is known to promote malignant transformation may be therapeutically intercepted before the tumor-prone region becomes cancerous.

Multiple studies have emphasized the essential role of *NOTCH3* in mediating heterotypic adhesion between endothelial and perivascular cells (27, 43, 44). We postulated various regulatory mechanisms for how perivascular cells might mediate the interaction of endothelial capillaries with the ECM. Integrin β1 has been shown to control VE-cadherin localization and blood vessel stability (45); however, silencing of the *ITGβ1* gene only exacerbated the dysregulation of vascular morphology and barrier function. TGFβ1 is a key regulator of pericyte differentiation (**Figure S19**) (46). Silencing of *TGFβ1* gene significantly reduced the negative effects of stiff bundled collagen on vascular capillaries; however, it also had seemingly negative effects on morphology and migration patterns in soft reticular collagen, which would be undesirable from a therapeutic perspective. Silencing of *NOTCH3*, on the other hand, was postulated to block the heterotypic cell communication between ECs and perivascular cells, thus preventing dysregulated perivascular cells from signaling endothelial cells from the damage on the abluminal ECM. Silencing *NOTCH3* had minimal visible effects on the soft reticular collagen vessels, which became leakier in sparse areas, as previously published (27). However, in stiff collagen the silencing resulted in the normalization of vasculature and a consequential reduction in IL-8 secretion (**Figure 5 A, E**). This points to a potential target in vascular therapies that may have important consequences. Blocking the mechano-paracrine feedback loop mediated by perivascular cells may reduce the pro-inflammatory responses and dysregulated remodeling of the vascular matrix, which may provide an effective therapeutic target for conditions in which fibrosis and inflammation are known causes of worse prognoses.

In conclusion, this study identified novel events and mechanisms of perivascular regulation of microvascular homeostasis in response to collagen stiffening and microarchitectural changes. Our results demonstrate, for the first time, that perivascular cells may be critical mediators of important events of matrix remodeling and perivascular inflammation that further exacerbate the effects of matrix fibrosis and chronic inflammation. We suggest that these findings may shine new light on the intersection of tissue fibrosis and vascular abnormalities that are common to many diseases, and may therefore have important implications for therapeutics, disease biology and regenerative medicine.

## Materials and Methods

### 1. Fabrication of the microfluidic device

A CAD program was used to design a mold comprised of two reservoirs connected by a central channel and a chamber. We then used a 3D printer (CADworks3D μMicrofluidic printer) and resin (Master Mold resin, CADworks) to print the positive molds. Printed micromolds were cleaned in methanol in three rinses of 2 min each under agitation, subsequently were cast with polydimethylsiloxane (PDMS, Sylgard 184, Dow-Corning), and left to cure overnight at 80°C, as previously described (22, 24). Next, PDMS was removed from the resin mold, two reservoirs were prepared using a 5-mm biopsy punch, for media, and 1-mm biopsy punches for the collagen loading ports. Molds were cleaned with ethanol, and immediately plasma bonded to a glass coverslip. Assembled devices were autoclaved, then treated with 1% (w/v) glutaraldehyde (Sigma) for 15 min, rinsed three times with distilled water (DIW), and left overnight in DI water to remove any trace of glutaraldehyde. To mold cylindrical channels, sterile 160-μm-diameter acupuncture needles were immersed in 0.1% BSA solution for at least 30 min, then inserted into the central channel of the devices approximately 200 μm above the glass coverslip surface. Rat tail collagen type I (3.0 mg/mL, BD Biosciences) was prepared according to the manufacturer’s protocol.

### 2. Collagen preparation

A stock solution of collagen type I (collagen I, rat tail Gibco, 3 mg/mL – cat #A10483-01) was prepared on ice with 10X PBS, 1N NaOH and endothelial growth media (EGM, Lonza) according to the manufacturer’s protocol. Briefly, to prepare 1 mL of a 2.5 mg/mL collagen solution, 833 µl of collagen was mixed with 100 µl of PBS, 20 µl of 1N NaOH and 47 µl of endothelial growth medium (EGM) and the working pH was 7.0-7.2. Thirty microliters of the final solution were immediately pipetted into the middle chamber, and allowed to polymerize at 37, 16, 21 or 4°C (23). To prevent collagen dehydration, after one hour, groups 37, 21 and 16°C had their main reservoirs filled with EGM, then devices were returned for incubation at their respective temperatures overnight. On the following day, cell medium was added to the reservoirs of the 4°C samples, allowed to incubate for 1 hour, then needles were carefully removed with a pair of tweezers, and cell medium (EGM) was replaced by fresh medium. Subsequently, the devices from all groups, with their reservoirs filled with fresh EGM, were placed in the incubator at 37°C, 5% CO2 overnight before cell seeding.

### 3. Scanning electron microscopy, quantification of collagen fiber thickness and pore area

Collagen scaffold structure and morphology were analyzed via scanning electron microscopy. To that end, 8 mm collagen disks (n = 3) were prepared as described above, cross-sectioned, fixed using a solution containing 2.5% glutaraldehyde in cacodylate buffer for two hours, dehydrated with graded ethanol solutions and lyophilized overnight. Samples were subsequently sputter-coated with gold/palladium and imaged using a FEI Helios Nanolab 660 DualBeam Scanning Electron Microscope at 20.0 kV or a FEI Quanta 200 SEM at 20.0 kV. For quantification of collagen fiber thickness and pore size, 3 ROI (left-center-right regions of interest) from each disk were imaged. Next, ImageJ was used to measure fiber thickness and pore area.

### 4. Atomic Force Microscopy

The collagen gels’ mechanical properties were measured by AFM, using a Bioscope Catalyst device (Bruker Santa Barbara, CA) attached to an inverted optical microscope (IX71, Olympus, Japan) in a similar manner as described previously (23). Briefly, the gels were probed under PBS with a V-shaped cantilever (conical tipped, nominal k=0.01N m^-1^ (Bruker, Santa Barbara, CA), whose spring constant was pre-calibrated by the thermal noise method in air. The relationship between photodiode signal and cantilever deflection was computed from the slope of the force displacement curve obtained at a bare region of a coverslip or cell culture dish. For each gel, we used force-volume mode in which we acquired multiple force-displacement (F-z) curves (where F=k * d, d being the deflection of the cantilever) by recording F and z while the piezo translator was ramped downward and upward at constant speed as it moved across an area as of 32 x 32 mm2 or 64 x 64 mm^2^ acquiring a force curve at every point of a predefined 32 x 32 array. We used 10– 100% of the baseline to trigger force fit boundaries for acquiring the Young’s modulus values. We only included data points having an R^2^ value greater than 0.9 and disregarded a small fraction of force curves with lower R^2^ values or those that did not show a predefined trigger force of typically 1 nN. To measure effective single fiber stiffness, the lower parts of the curves (typically, up to 10-20 nm indentation) were fitted by a linear function whose slope is the fiber stiffness.

### 5. Cell culture

Human umbilical vein endothelial cells expressing green fluorescent protein (GFP-HUVECs) (C2519A, Lonza, Basel, Switzerland) were cultured in a supplemented (EGM-2 bullet kit, Lonza) endothelial cell growth medium (Lifeline Cell Technology, California, USA). Human bone marrow mesenchymal stem cells (hMSCs, RoosterBio, Maryland, USA) were cultured in alpha Minimal Eagle’s medium (α-MEM) (Gibco, Carlsbad, USA) with 10% FBS (cat# 16141079, Gibco) and 1% penicillin/streptomycin (cat # 15140122). Cell media were changed every other day, and cells were passaged when reaching a confluency of 80–90%. HUVECS at passages 4–6 and hMSCs at passages 2–4 were used for all the experiments. All cells were maintained in a humidified incubator (5% CO2, 37°C).

### 6. Cell seeding

For seeding, GFP-HUVECs and hMSCs were trypsinized, counted and mixed in 4:1 ratio according to previous publications (47, 48) in a cell density of 6 million cells/mL. Subsequently, the cell medium was removed from the reservoirs, and 25 µl of the cell suspension was added into one reservoir. The devices were flipped upside down, placed in the incubator for 5 min, seeded again and left upside down in the incubator for another 5 min. Until the entire extension of the collagen channel had cells attached, we repeated the seeding, flipping the chip as needed. Next, the devices were placed in the incubator for 30 min under static conditions. Afterwards, the devices were transferred to the 2D rocker (BenchRocker) inside the incubator, as published previously (22, 24).

### 7. Cell staining and imaging

After 48h in culture, samples were rinsed with phosphate-buffered saline (PBS), fixed with buffered formalin 10% in PBS (v/v) for 30 min, washed with PBS, permeabilized with 0.1% (w/v) Triton X-100 for 10 min, and blocked with 1.5 % (w/v) bovine serum albumin (BSA) for 1 h under agitation. After washing with PBS, samples were incubated with one of the following primary antibodies (anti-PECAM-1, rabbit anti-human, cat#AB32457, Abcam, 1:100; anti-N-cadherin, rabbit anti-human, cat#AB18203, Abcam, 1:100; anti-laminin, rabbit anti-human, cat#PA1-16730, Invitrogen, 1:200; anti-NG-2, mouse anti-human, cat#14-6504-82, Invitrogen, 1:200; anti-phosphorilated paxillin, rabbit anti-human, cat#NBP2-81063-UV, Novus Biologicals) overnight at 4°C. Samples were washed with PBS and incubated with secondary antibody (goat anti-mouse Alexa Fluor 555, cat#A21422Invitrogen, 1:250; goat anti-rabbit Alexa Fluor 647, cat#21244, Invitrogen, 1:250) for overnight at 4°C under agitation. This was followed by rinsing in 0.1% PBS, staining of the nuclei using NucBlue (Fixed Cell ReadyProbes, DAPI, cat#R37606, Molecular Probes) and staining actin with ActinGreen 488 (ReadyProbes, cat#R37110, Molecular Probes) for 1 h at 37 °C under agitation.

Samples were imaged using a confocal microscope (Zeiss, LSM 880, Germany) with a 10× objective (N.A. 0.45, Zeiss, Plan Apochromat). The depth of imaging was 100–400 μm, split in at least 100 Z-stacks. Z-stacks were converted into TIFF files or 3D images using Zen or Imaris software (v9.1, Bitplane, Oxford Instruments, Zurich, Switzerland). To quantify sprout density and length in the scaffolds, 3 images of each device were obtained. Image J and Imaris were used to measure the individual distances from the leading protrusions of tip cells to the wall of the parent vessel (n=3 samples per condition, 3 ROI per sample). Migrating cells were quantified using the plugin ‘Cell counter’ from ImageJ: only cells that had no contact with the parent vessel and did not stain for PECAM-1 were considered (n=3 samples per condition, 3 ROI per sample). To quantify NG2+cells associated with the capillaries the images were opened in ImageJ and only cells that were in contact with the endothelial cells were considered. For PECAM-1 and phosphorylated paxillin quantification, images were converted to binary, skeletonized in ImageJ, and normalized by the number of nuclei. Laminin was measured with ImageJ as a function of vessel area.

### 8. Dextran Assay for Barrier Function Measurement

To measure the permeability of the endothelium in the microfluidic platform, fluorescent dextran (10 kDa Cascade Blue & 70 kDa Texas Red, Thermo Fisher) was introduced into perfusion media (EGM2) at a concentration of 12.5 μg/mL. Diffusion of the dextran was imaged in real time with a confocal microscope (LSM 880, Carl Zeiss) at 10x magnification. The diffusive permeability coefficient (Pd) was calculated by measuring the flux of dextran into the collagen gel and fitting the resulting diffusion profiles to a dynamic mass conservation equation as described previously (15). Next, 70 kDa dextran was perfused through the vessel lumens and extravasation of the dextran was measured as a function of time to quantify the diffusive permeability.

### 9. Nanostring and Differential Expression Analysis

Single channels within the collagen (∼150,000 cells) were removed from each device, then suspended in 0.05 ml in RIPA buffer (ThermoFisher) at 4°C, vortexed for 10 min at 4 °C, lysed and RNA content was measured using a nanodrop. Only samples with at least 100 μg of RNA content per μl with RLT buffer (Qiagen) + 1% 2-mercaptoethanol (Sigma) were utilized. A volume of 1.5 μl of lysates were used in the nanoString hybridization reaction and the remainder was stored at −80 °C. nCounter Elements (nanoString) hybridization was performed according to the manufacturer’s instructions. The Pan Cancer Progression panel was used and the list of gene targets is given in the **Dataset S1**.

Differential expression analysis was performed using the R package DESeq2 (v 1.24.0). The filtered counts table was normalized using DESeq2’s median of ratios method of normalization. Differentially expressed genes were then identified using the negative binomial GLM method in DESeq2 with a p-value cutoff of 0.01 and a fold change cutoff of 2. The differential gene list was then sorted by adjusted p-value and the top 50 genes were plotted on a heatmap using pheatmap (v1.0.12). The full differential expression results were also plotted on an MD plot and a volcano plot using the functions glMDPlot and glXYPlot from the package glimma (v1.10.1). The pathways and enrichment scores were obtained using David Annotation tool and plotted with SRPLOT.

### 10. siRNA transfection

hMSCs were plated into a 6-well plate in antibiotic-free alphaMEM (Gibco) 24 h before siRNA transfection. Transfection reagents and siRNA for NOTCH3, TGFB1, IntegrinB1 and ON-Target plus cyclophilin B control pool (ON-TARGET plus siRNA kits, Horizon Discovery - cat # L-011093-00-0005, L-012562-00-0005, L-004506-00-0005, D-001820-10-05) were diluted in Opti-MEM and added to the wells of 80% confluent cells, according to manufacturer instructions (27). Each gene was silenced in different well plates to avoid any cross contamination during the procedures. Next, 2 μL of transfection reagent was used per well, and siRNA were added for a final concentration of 40 nM. After 24 h, cells were passaged and used in downstream experiments. Cells were lysed for Western blot at 48 h post-transfection to quantify knockdown. This timepoint coincided with the end point of the single channel experiments. Three independent experiments were performed and the graphics represent quantification of these independent samples.

### 11. Engineering bone marrow, seeding on the vascularized chip and flow cytometry

Bone marrow organoids were fabricated by seeding 3 × 10^3^ mesenchymal stem cells (hMSC, Lonza, PT-2501) and 6 × 10^3^ human bone marrow CD34+ cells (STEMCELL Technologies, 70002.3) into a collagen-based hydrogel according to the protocol described previously (49) . These samples were cultured for seven days in StemSpan SFEM II (STEMCELL Technologies, 09655) supplemented with a cytokine cocktail that supports hematopoietic stem cell growth and differentiation, the basis of which is described in Chou et al. 2020 (49). After seven days of culture under standard conditions, cells were harvested from the three-dimensional matrix using Collagenase Type I (Gibco, 17018-029) and NSK Fermented Soybean Extract (Japan Bio Science Laboratory Co., Ltd., NSK-SD). A total of 6 × 10^3^ cells was reseeded into an identical hydrogel.

Next, microfluidic devices for the single channels had the top of the collagen chamber removed with a 6-mm biopsy punch, then hollow channels were fabricated with acupuncture needles and soft reticular or stiff bundled collagen as described above. GFP-HUVECs and hMSCs were seeded and after 24 h, bone-marrow loaded hydrogel was overlaid on the vascularized chip inlet well, and solidified at 37°C (Supplementary Figure 20). The chip was cultured for 72 hours on a 2D rocker under standard conditions in the supplemented StemSpan SFEM II media. Cell media were changed every day.

Prior to flow cytometry, the hydrogels were digested with Collagenase Type I (Gibco, 17018-029) and NSK Fermented Soybean Extract (Japan Bio Science Laboratory Co., Ltd., NSK-SD). The cells were stained with Live/Dead Green Fixable Dead Cell Stain (Invitrogen, L34970), PE Mouse Anti-Human CD16 (3G8, BD Biosciences), BV421 Mouse Anti-Human CD33 (WM53, BD Biosciences), APC Mouse Anti-Human CD34 (581, BD Biosciences), PerCP-Cy™5.5 Mouse Anti-Human CD45 (HI30, BD Biosciences), APC/Cyanine7 anti-human CD90 (5E10, BioLegend), PE-Cy™5 Mouse Anti-Human CD235a (HIR2, BD Biosciences), APC-R700 Mouse Anti-Human CD117 (YB5.B8, BD Biosciences), and BUV805 Mouse Anti-Human CD14 (MφP9, BD Biosciences). Stained cells were washed and resuspended with CountBright Absolute Counting Beads (Invitrogen, C36950) and Flow Cytometry Staining Buffer (Invitrogen eBioscience, 00422226). Data were collected using a Cytek Aurora 5-laser spectral cytometer and SpectroFlo software (Cytek). The unmixed data were gated and analyzed with FlowJo v10.9.0. The cells of interest were isolated by gating out debris, doublets and dead cells.

Monocytes were identified as CD45High CD33High CD34-CD117-CD16-CD14+. Neutrophils were identified as CD45High CD14-CD34-CD33+/-CD16+. Mesenchymal stem cells were identified as CD45Low CD235a-CD34-CD90+ cells. The distinction between CD33High and CD33Low cells was made downstream of CD45High.

### 12. ELISA

The IL-8 and TGF-β1 production was measured from the supernatants of three biological replicate chips from soft reticular and stiff bundled groups after 48 h of incubation. The concentration of both cytokines was determined by a RayBio® Human IL-8 ELISA Kit (RayBiotech, Peachtree Corners, GA) and RayBio® Human TGFβ1 ELISA Kit (RayBiotech, Peachtree Corners, GA) following the manufacturer’s instructions. The results were compared to a colorimetric standard curve (4000-16.38 pg/mL for TGFβ1 and 600-0.8 pg/mL) after reading the microplates in a microplate reader (SPARK 20M, TECAN, Seestrasse, Männedorf, Switzerland).

### 13. Statistical analysis

All experiments were done at least in triplicate. Data are presented as mean ± SD, statistical analyses were performed using GraphPad Prism (version 9, GraphPad Software, LLC) using one-way or two-way ANOVA and Tukey’s post hoc tests (α=0.05). For pairwise comparisons Student’s t-test was used. Statistically significant differences were determined as *(*P* < 0.05), **(*P* < 0.01), ***(*P* < 0.001), and ****(*P* < 0.0001) respectively.

## Acknowledgments

This project was supported by funding from the National Institute of Dental and Craniofacial Research (R01DE026170, R01DE026170-03S1, R01DE026170-03S2, R01DE029553 to L.E.B., and K01DE030484-01A1 to CMF, PORT T90 fellowship to A.A.), CAPES/Brazil# 88887.716956/2022-00 to ACSS. M.U. was supported by an EMBO long-term fellowship (EMBO ALTF811-2018), the Center for Multiscale & Translational Mechanobiology at Boston University, and an AHA postdoctoral fellowship (828475). K.M.Y. was supported by ZIADE000718 and ZIADE000719. The Advanced Light Microscopy Core was supported by the National Cancer Institute - P30CA069533. K.B and M.F are funded by the Cancer Early Detection Advanced Research Center (CEDAR). We acknowledge Jinho Lee, from the Nanostring Laboratory at OHSU Knight Diagnostic Laboratories for assisting in performing the Nanostring assay. We extend our acknowledgment to the Hugo Cross and Theresa Lusardi for their aid in gene analyses, Sean D. Speese and Stefanie Kaech Petrie for their valuable expertise in confocal imaging and image analysis.

## Notes

### Competing Interest Statement

The authors have declared no competing interest.

## References

1. P. Carmeliet, R. K. Jain, Molecular mechanisms and clinical applications of angiogenesis. Nature 473, 298–307 (2011).

2. A. Armulik, G. Genove, C. Betsholtz, Pericytes: developmental, physiological, and pathological perspectives, problems, and promises. Dev Cell 21, 193–215 (2011).

3. D. von Tell, A. Armulik, C. Betsholtz, Pericytes and vascular stability. Exp Cell Res 312, 623–629 (2006).

4. A. J. Engler, S. Sen, H. L. Sweeney, D. E. Discher, Matrix elasticity directs stem cell lineage specification. Cell 126, 677–689 (2006).

5. D. E. Discher, P. Janmey, Y. L. Wang, Tissue cells feel and respond to the stiffness of their substrate. Science 310, 1139–1143 (2005).

6. M. G. Jones et al., Nanoscale dysregulation of collagen structure-function disrupts mechano-homeostasis and mediates pulmonary fibrosis. Elife 7 (2018).

7. C. Lyu et al., Advanced glycation end-products as mediators of the aberrant crosslinking of extracellular matrix in scarred liver tissue. Nat Biomed Eng 7, 1437–1454 (2023).

8. B. R. Seo et al., Collagen microarchitecture mechanically controls myofibroblast differentiation. Proc Natl Acad Sci U S A 117, 11387–11398 (2020).

9. M. J. Paszek et al., Tensional homeostasis and the malignant phenotype. Cancer Cell 8, 241–254 (2005).

10. C. A. Dessalles, C. Leclech, A. Castagnino, A. I. Barakat, Integration of substrate- and flow-derived stresses in endothelial cell mechanobiology. Commun Biol 4, 764 (2021).

11. N. Monteiro, W. He, C. M. Franca, A. Athirasala, L. E. Bertassoni, Engineering Microvascular Networks in LED Light-Cured Cell-Laden Hydrogels. ACS Biomater Sci Eng 4, 2563–2570 (2018).

12. C. M. Nelson, D. M. Pirone, J. L. Tan, C. S. Chen, Vascular endothelial-cadherin regulates cytoskeletal tension, cell spreading, and focal adhesions by stimulating RhoA. Mol Biol Cell 15, 2943–2953 (2004).

13. J. C. Kohn et al., Cooperative effects of matrix stiffness and fluid shear stress on endothelial cell behavior. Biophys J 108, 471–478 (2015).

14. N. Koike et al., Tissue engineering: creation of long-lasting blood vessels. Nature 428, 138–139 (2004).

15. S. Alimperti et al., Three-dimensional biomimetic vascular model reveals a RhoA, Rac1, and. *Proc Natl Acad Sci U S A* **114**, 8758-8763 (2017).

16. J. Kim et al., Engineering of a Biomimetic Pericyte-Covered 3D Microvascular Network. PLoS One 10, e0133880 (2015).

17. S. Dulauroy, S. E. Di Carlo, F. Langa, G. Eberl, L. Peduto, Lineage tracing and genetic ablation of ADAM12(+) perivascular cells identify a major source of profibrotic cells during acute tissue injury. Nat Med 18, 1262–1270 (2012).

18. K. Stark et al., Capillary and arteriolar pericytes attract innate leukocytes exiting through venules and ’instruct’ them with pattern-recognition and motility programs. Nat Immunol 14, 41–51 (2013).

19. M. Murgai et al., KLF4-dependent perivascular cell plasticity mediates pre-metastatic niche formation and metastasis. Nat Med 23, 1176–1190 (2017).

20. Y. Yang et al., The PDGF-BB-SOX7 axis-modulated IL-33 in pericytes and stromal cells promotes metastasis through tumour-associated macrophages. Nat Commun 7, 11385 (2016).

21. L. Díaz-Flores et al., Pericytes. Morphofunction, interactions and pathology in a quiescent and activated mesenchymal cell niche. Histol Histopathol 24, 909–969 (2009).

22. W. J. Polacheck, M. L. Kutys, J. B. Tefft, C. S. Chen, Microfabricated blood vessels for modeling the vascular transport barrier. Nat Protoc 14, 1425–1454 (2019).

23. A. D. Doyle, N. Carvajal, A. Jin, K. Matsumoto, K. M. Yamada, Local 3D matrix microenvironment regulates cell migration through spatiotemporal dynamics of contractility-dependent adhesions. Nat Commun 6, 8720 (2015).

24. D. H. Nguyen et al., Biomimetic model to reconstitute angiogenic sprouting morphogenesis in vitro. Proc Natl Acad Sci U S A 110, 6712–6717 (2013).

25. K. C. P. Cheung et al., Preservation of microvascular barrier function requires CD31 receptor-induced metabolic reprogramming. Nat Commun 11, 3595 (2020).

26. A. M. López-Colomé, I. Lee-Rivera, R. Benavides-Hidalgo, E. López, Paxillin: a crossroad in pathological cell migration. J Hematol Oncol 10, 50 (2017).

27. J. B. Tefft et al., Notch1 and Notch3 coordinate for pericyte-induced stabilization of vasculature. Am J Physiol Cell Physiol 322, C185–C196 (2022).

28. Y. Yu et al., Extracellular Matrix Stiffness Regulates Microvascular Stability by Controlling Endothelial Paracrine Signaling to Determine Pericyte Fate. Arterioscler Thromb Vasc Biol 43, 1887–1899 (2023).

29. P. Navarro, L. Ruco, E. Dejana, Differential localization of VE- and N-cadherins in human endothelial cells: VE-cadherin competes with N-cadherin for junctional localization. J Cell Biol 140, 1475–1484 (1998).

30. K. L. Mui et al., N-Cadherin Induction by ECM Stiffness and FAK Overrides the Spreading Requirement for Proliferation of Vascular Smooth Muscle Cells. Cell Rep 10, 1477–1486 (2015).

31. J. D. Baranski et al., Geometric control of vascular networks to enhance engineered tissue integration and function. Proc Natl Acad Sci U S A 110, 7586–7591 (2013).

32. A. J. Dunbar et al., CXCL8/CXCR2 signaling mediates bone marrow fibrosis and is a therapeutic target in myelofibrosis. Blood 141, 2508–2519 (2023).

33. E. Shimshoni et al., Distinct extracellular-matrix remodeling events precede symptoms of inflammation. Matrix Biol 96, 47–68 (2021).

34. K. G. Danielson et al., Targeted disruption of decorin leads to abnormal collagen fibril morphology and skin fragility. J Cell Biol 136, 729–743 (1997).

35. T. Hashimoto et al., Inhibition of MDM2 attenuates neointimal hyperplasia via suppression of vascular proliferation and inflammation. Cardiovasc Res 91, 711–719 (2011).

36. M. Farhang Ghahremani et al., p53 promotes VEGF expression and angiogenesis in the absence of an intact p21-Rb pathway. Cell Death Differ 20, 888–897 (2013).

37. X. Liu et al., Niche stiffness sustains cancer stemness via TAZ and NANOG phase separation. Nat Commun 14, 238 (2023).

38. W. Wang et al., Diabetic hyperglycemia promotes primary tumor progression through glycation-induced tumor extracellular matrix stiffening. Sci Adv 8, eabo1673 (2022).

39. T. B. Cruz et al., Mice with Type 2 Diabetes Present Significant Alterations in Their Tissue Biomechanical Properties and Histological Features. Biomedicines 10 (2021).

40. R. Chen, D. G. McVey, D. Shen, X. Huang, S. Ye, Phenotypic Switching of Vascular Smooth Muscle Cells in Atherosclerosis. J Am Heart Assoc 12, e031121 (2023).

41. H. Y. Tang et al., Vascular Smooth Muscle Cells Phenotypic Switching in Cardiovascular Diseases. Cells 11 (2022).

42. N. Bansaccal et al., The extracellular matrix dictates regional competence for tumour initiation. Nature 623, 828–835 (2023).

43. K. Gaengel, G. Genove, A. Armulik, C. Betsholtz, Endothelial-mural cell signaling in vascular development and angiogenesis. Arterioscler Thromb Vasc Biol 29, 630–638 (2009).

44. H. Zeng et al., LPS causes pericyte loss and microvascular dysfunction via disruption of Sirt3/angiopoietins/Tie-2 and HIF-2alpha/Notch3 pathways. Sci Rep 6, 20931 (2016).

45. H. Yamamoto et al., Integrin β1 controls VE-cadherin localization and blood vessel stability. Nat Commun 6, 6429 (2015).

46. T. Yamazaki et al., Tissue Myeloid Progenitors Differentiate into Pericytes through TGF-β Signaling in Developing Skin Vasculature. Cell Rep 18, 2991–3004 (2017).

47. S. P. Parthiban et al., Engineering pericyte-supported microvascular capillaries in cell-laden hydrogels using stem cells from the bone marrow, dental pulp and dental apical papilla. Sci Rep 10, 21579 (2020).

48. C. M. Franca et al., The influence of osteopontin-guided collagen intrafibrillar mineralization on pericyte differentiation and vascularization of engineered bone scaffolds. J Biomed Mater Res B Appl Biomater (2018).

49. D. B. Chou et al., On-chip recapitulation of clinical bone marrow toxicities and patient-specific pathophysiology. Nat Biomed Eng 4, 394–406 (2020).

