## Supporting Information for "Perivascular cells function as mechano-structural sensors of vascular capillaries"

* Luiz E. Bertassoni

**This PDF file includes:**

Supporting text

Figures S1 to S21

**Other supporting materials for this manuscript include the following:**

Movies S1 to S10

Dataset S1

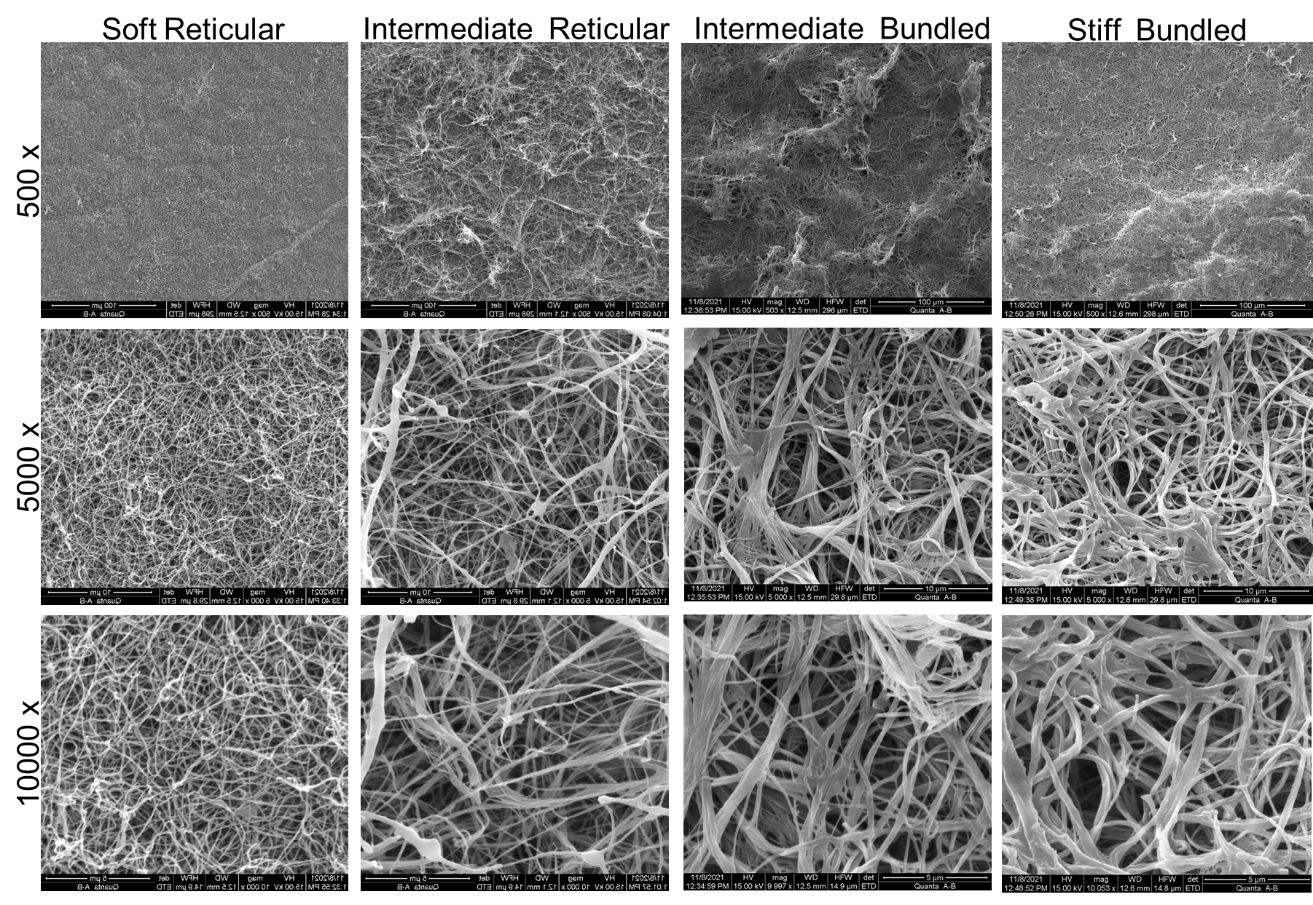

Figure S1. SEM of collagen microarchitectures as a function of fibrillogenesis temperature. From left to right – soft reticular (37^o^C), intermediate reticular (21^o^C), intermediate bundled (16^o^C) and stiff bundled (4^o^C). Scanning electron microscopy images show that polymerization at higher temperatures results in a soft reticular collagen with delicate fibers and homogeneous pore distribution. As polymerization temperature is decreased, fiber thickness and pore sizes increase in a controllable manner, progressively bundling collagen fibers without changing collagen density and the presence of cell ligands.

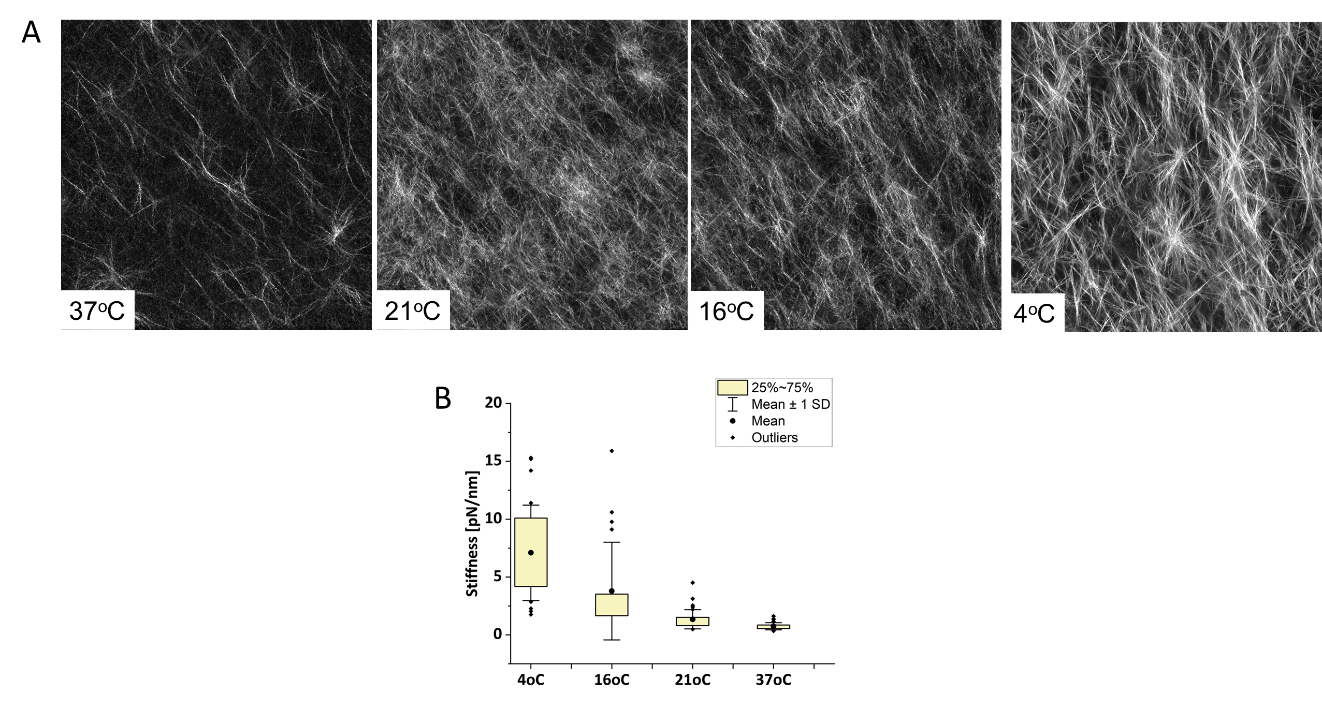

Figure S2. Atomic force microscopy images and measurements of collagen polymerized at different temperatures. (A) Collagen fibers became progressively thicker and more bundled with lower temperatures in a controllable manner. (B) According to AFM measurements, collagen polymerized at 37^o^C presents single fiber stiffness at an average range of 0.73±0.3 pN/nmm with the least variability. Decreasing polymerizing temperatures correlate with incremental increases in single fiber stiffness and greater variability among the fibers.

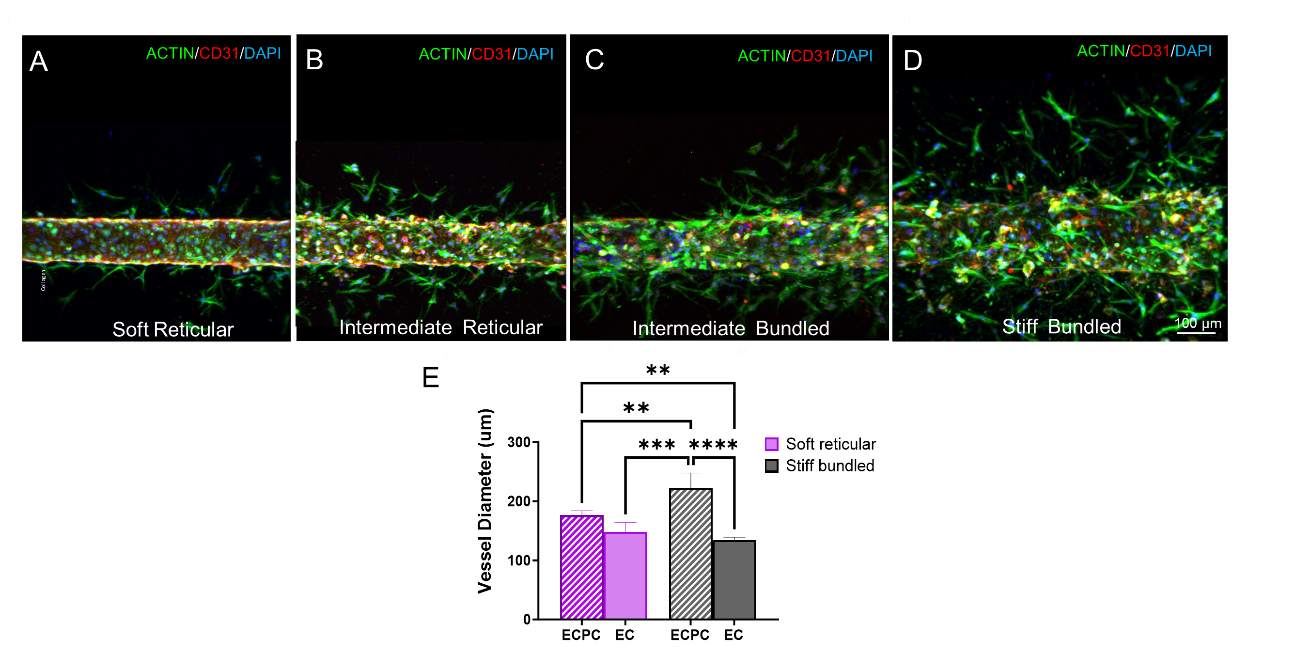

**Figure S3. Progressive changes in vascular morphology as a function of collagen stiffness and microarchitecture.** (A-D) Images depicting vasculature engineered with endothelial and perivascular cells showing a progressive alteration in vascular morphology and cell migration in relation to varying collagen stiffness and microarchitecture. (B) The association of perivascular cells with stiff bundled collagen results in an increased vessel diameter. In contrast, vasculature constructed solely with endothelial cells exhibited consistent diameter regardless of the collagen's composition.

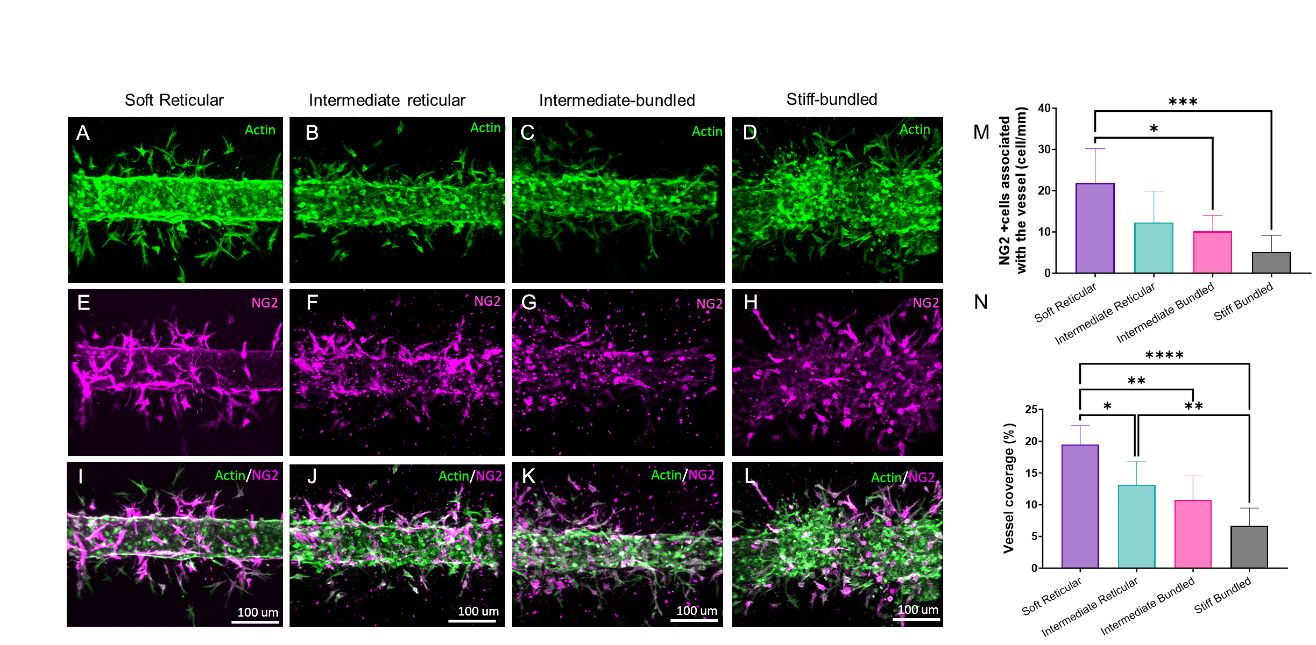

**Figure S4**. **Pericyte coverage as a function of collagen stiffness and microarchitecture**. Representative images depicting a progressive reduction in NG2 (pericyte marker) across varying collagen stiffness levels. (A,E,I) Soft reticular collagen resulted in NG2+ cells associated with the endothelial cells (D,H,L). On the other hand, in stiff bundled collagen, NG2+ cells migrated further from the capillaries, which are not characterized as pericytes due to their lack of contact with endothelial cells. Intermediate states (B,C,F,G,J,K) showed a progressive increase and migration of NG2+ cells away from the vasculature. (M,N) Quantitative analysis revealed a marked increase in pericyte differentiation within soft reticular collagen. Specifically, only NG2+ cells proximal to the vascular wall were identified as functional pericytes providing vessel coverage, with capillaries engineered within softer collagen exhibiting the highest pericyte coverage.

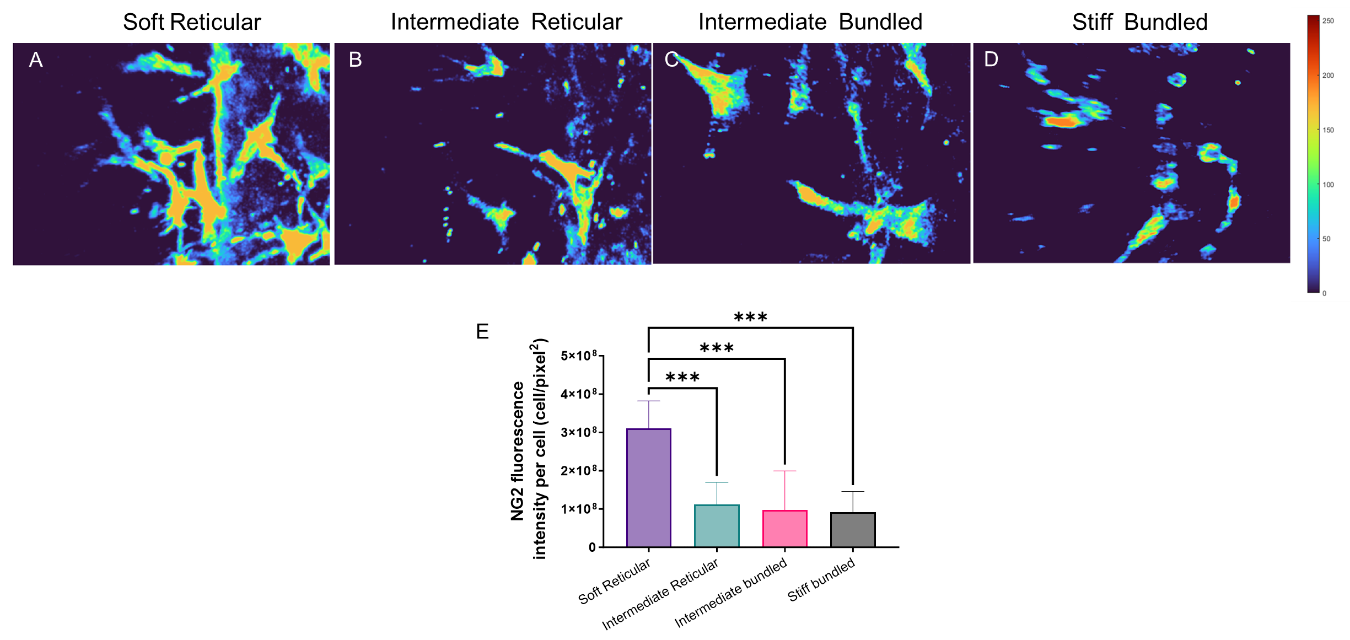

**Figure S5**. **Intensity of NG2 expression**. (A) In addition to a higher number of perivascular cells expressing NG2, vascular capillaries engineered in soft reticular collagen also demonstrated a higher intensity of NG2 expression per cell than (B,C) intermediate states or (D) stiff bundled collagen. (E) Quantification was performed to count cells that were associated with the capillaries.

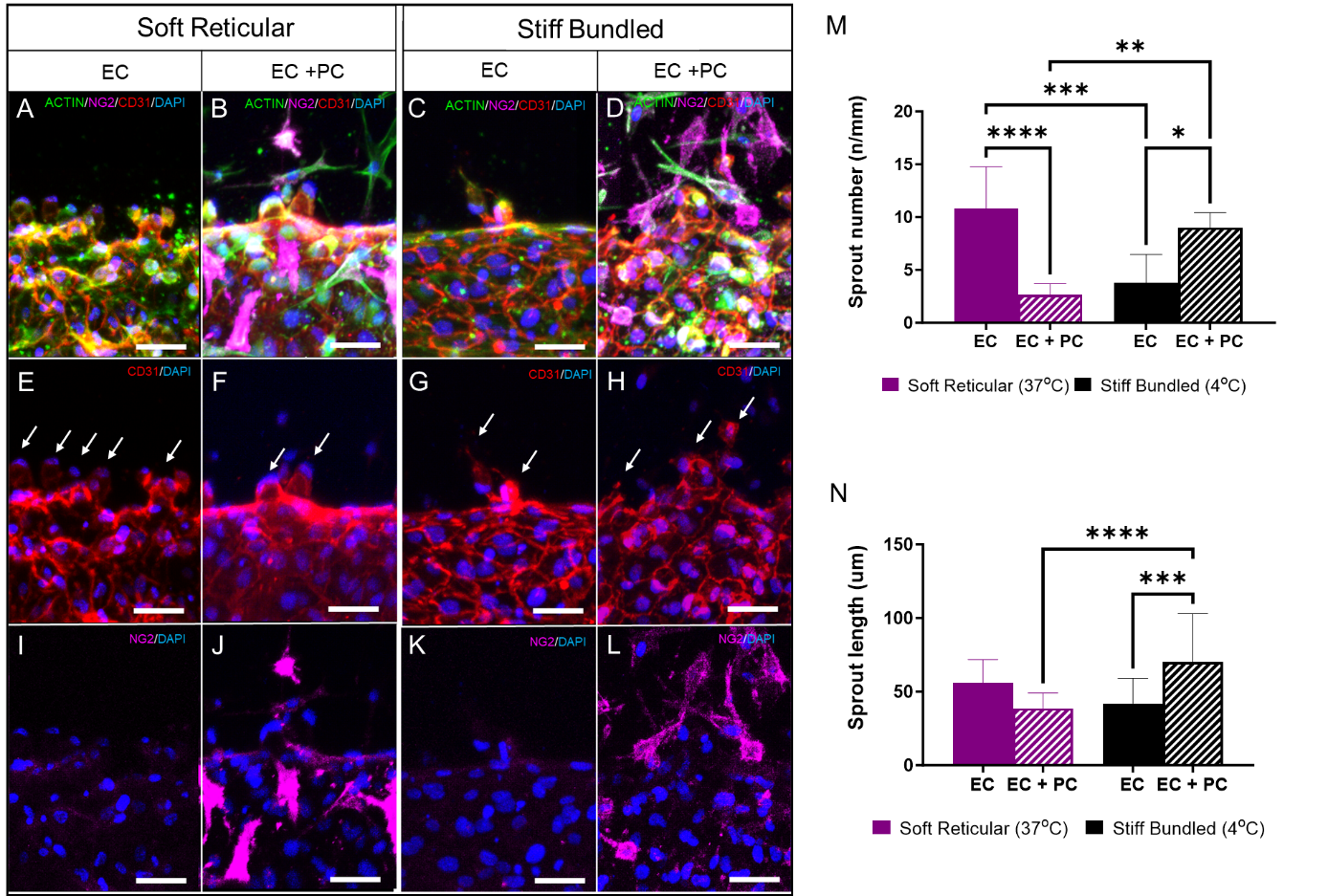

**Figure S6**. **Sprouts according to the presence of perivascular cells and collagen stiffness**. Comparative analysis of sprouting angiogenesis (arrows) reveals distinct outcomes in collagen matrices based on the presence or absence of perivascular cells (PC). (A,E,I, C,G,K, M,N) The absence of PC exhibited a higher sprout count in soft (healthy) collagen matrices compared to stiff (fibrotic) ones, while sprout length remained consistent across all groups. In contrast, the presence of PC (B,F,J,D,H,L, M,N) correlated with increased sprout number and length specifically within the stiff group, indicating differential responses in the presence of perivascular cells.

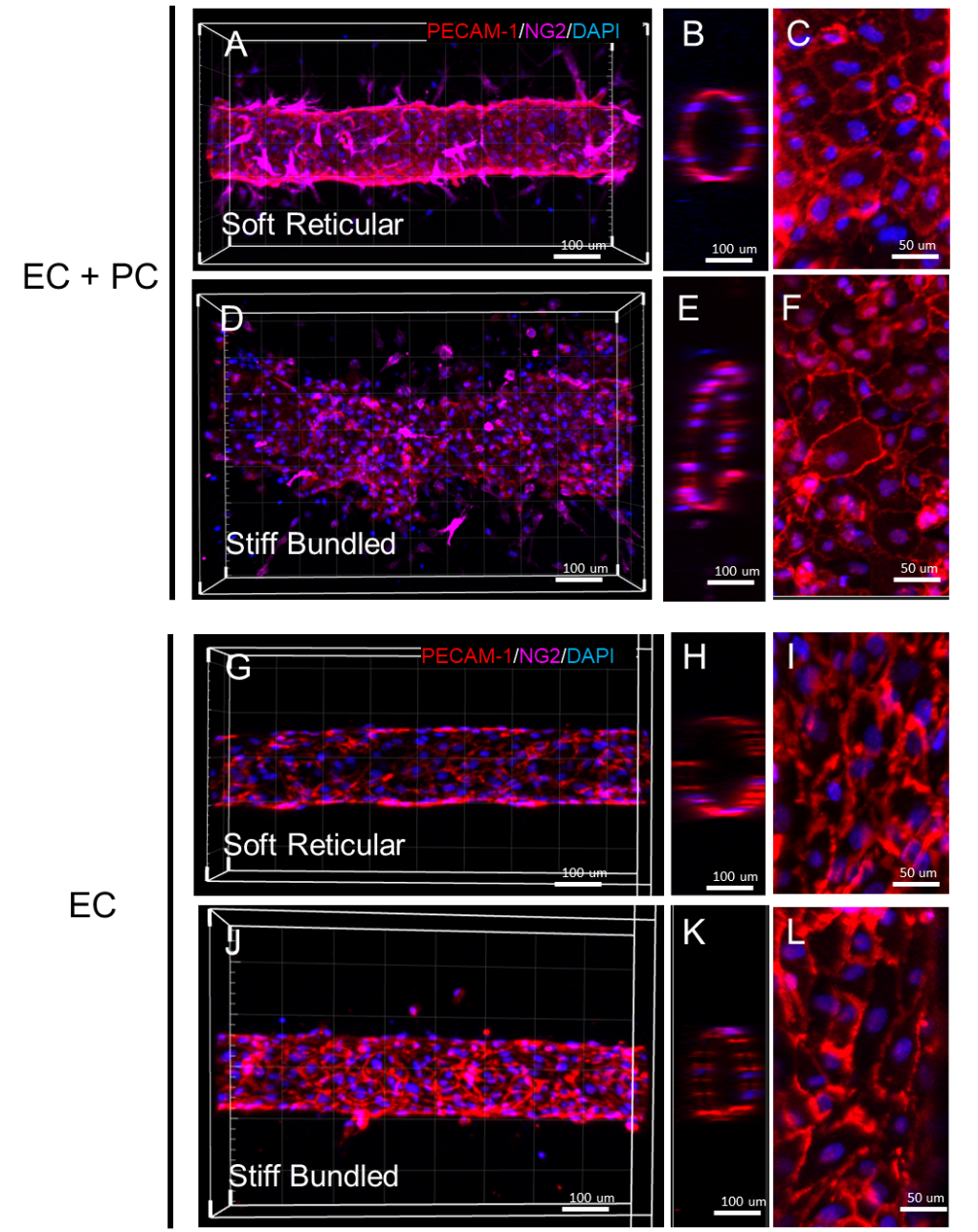

**Figure S7. PECAM-1 expression in capillaries engineered within soft reticular or fibrillar bundled collagen as a function of the presence of perivascular cells**. (A-C) Endothelial cells size was more homogeneous in the soft reticular collagen (healthy) in the presence of perivascular cells. (D-F) Stiff bundled collagen (more fibrotic) and perivascular cells led to irregular cell morphology with variable size and PECAM-1 expression. In contrast, when the vasculature was engineered with endothelial cells alone, the differences between cells in the two collagen microenvironments were mitigated (G-L)

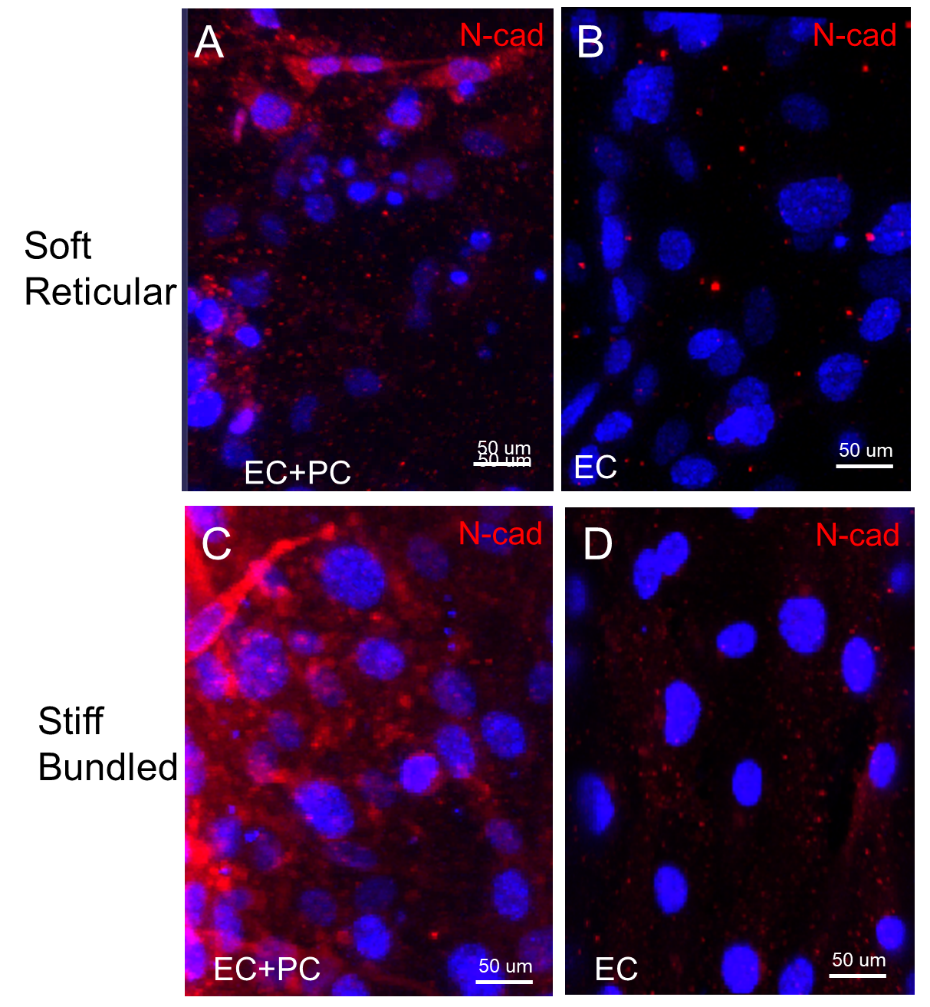

**Figure S8. Immunofluorescence analysis depicting N-cadherin expression patterns as a function of perivascular cells and collagen status**. N-cadherin expression was prominently observed in the stiff bundled collagen group with perivascular cells, while in the soft reticular group, it was predominantly confined to perivascular cells. Notably, in the absence of perivascular cells, endothelial cells in the stiff bundled group displayed limited N-cadherin expression.

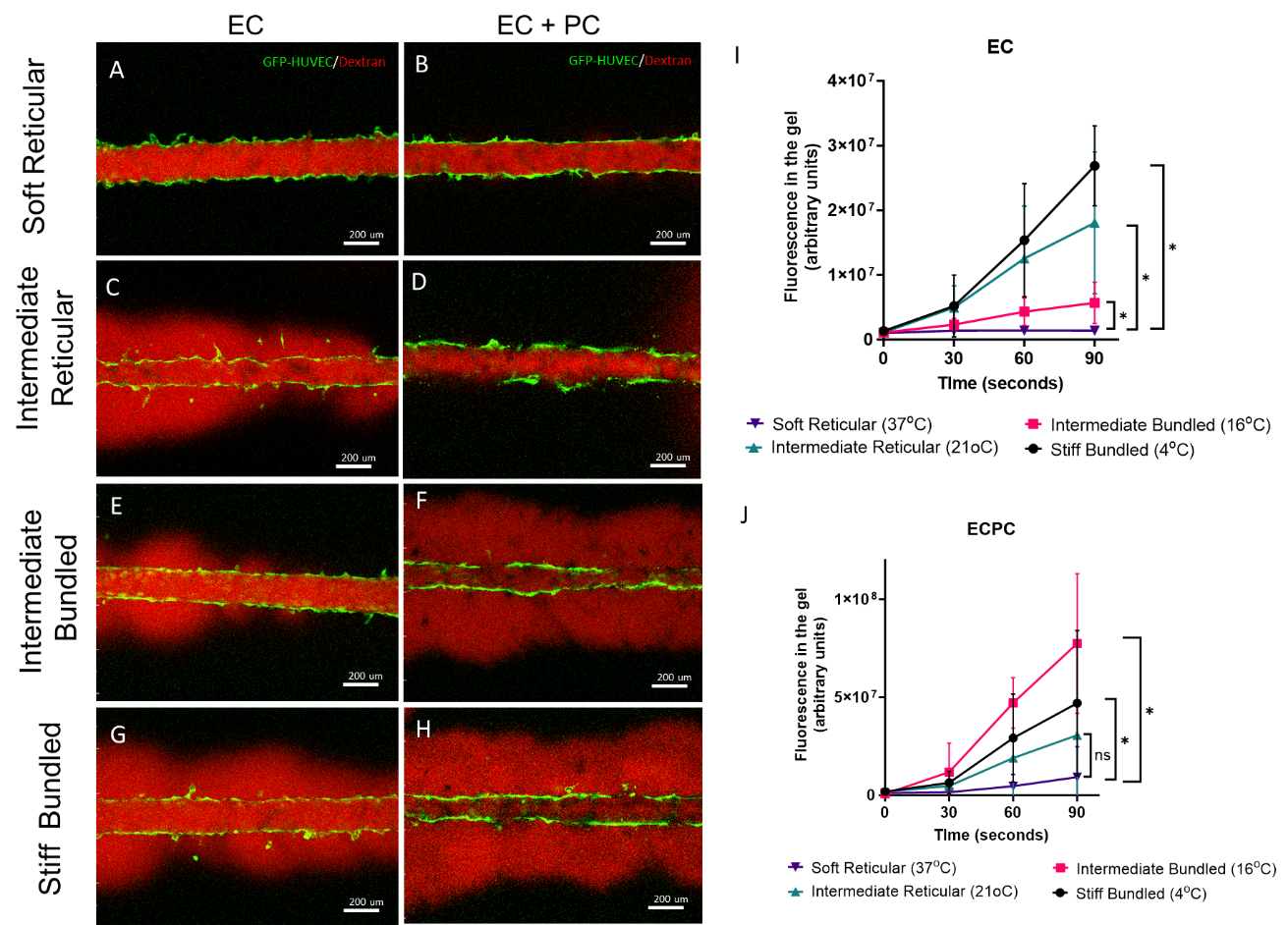

**Figure S9.** **Barrier function associated with perivascular cells in different collagen stiffness and architecture** – Representative images demonstrating dextran assay results after perfusion. Preservation of barrier function was evident in soft reticular collagen, irrespective of perivascular cell presence (A,B). In contrast, the absence of perivascular cells led to the loss of barrier function in all other groups without PC (C,E,G) and in the intermediate and stiff bundled groups with PC (F,H). Notably, perivascular cells exhibited sustained barrier function, even with a minor stiffness increment (D). However, in collagen configurations with increased stiffness and bundling, the presence of perivascular cells proved insufficient to maintain barrier function (I,J).

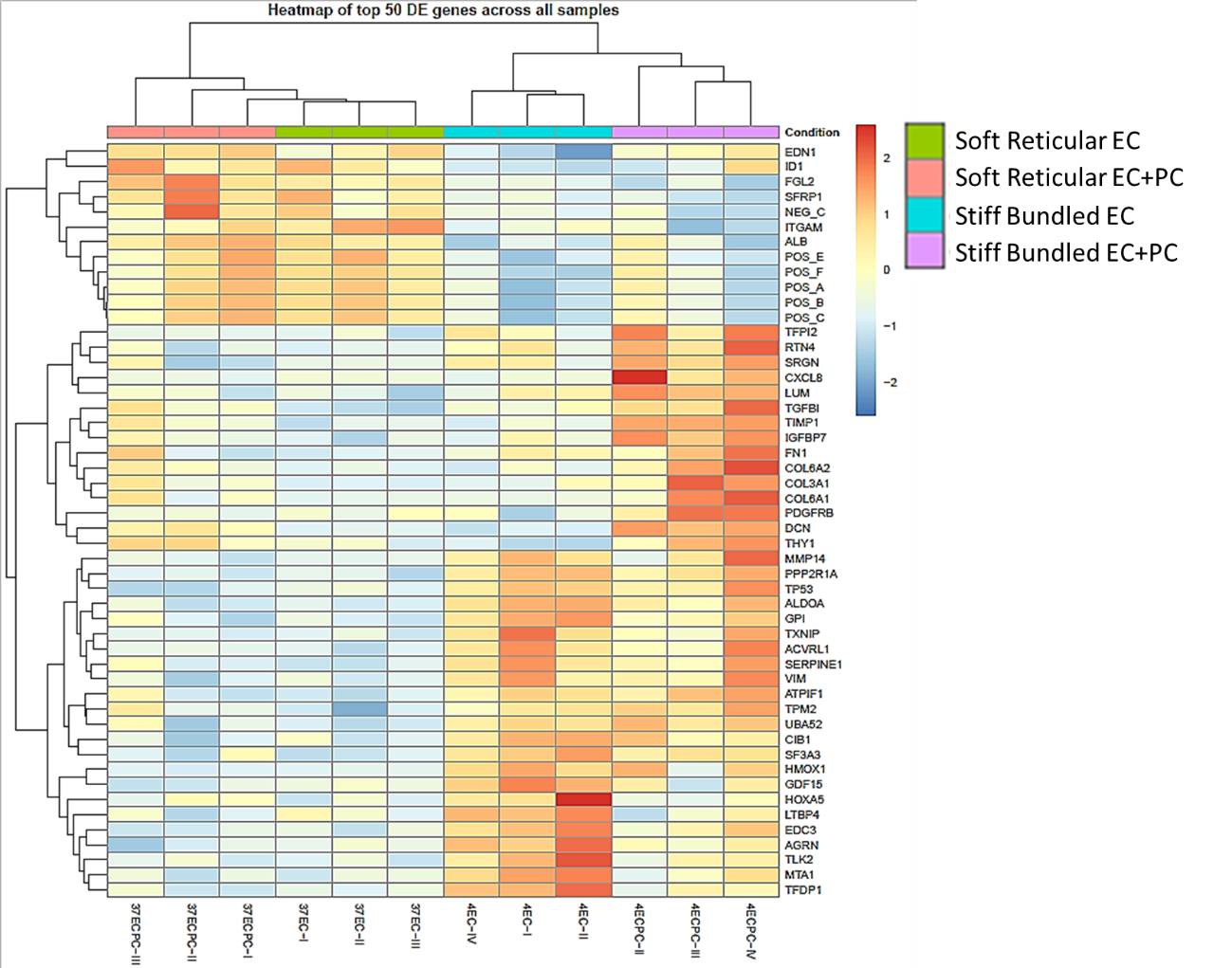

**Figure S10**. **Heat map** **highlighting the 50 most differentially expressed genes in stiff (more fibrotic) and soft (healthy) collagen vasculature with and without perivascular cells.** Analysis of gene expression profiles demonstrates distinct patterns in different collagen environments. Stiff collagen with perivascular cells exhibits upregulation of genes including *CXCL8, FN1, DCN, COL3A1, COL6A1, MMP14*, and *PDGFRb5*. Conversely, in the absence of perivascular cells, stiff collagen shows increased expression of *AGRN, MTA, TFDP*, and *HOXA5*. Soft collagen, on the other hand, displays a smaller subset of differentially expressed genes, including *ID1, FGL2,* and *ITGAM*.

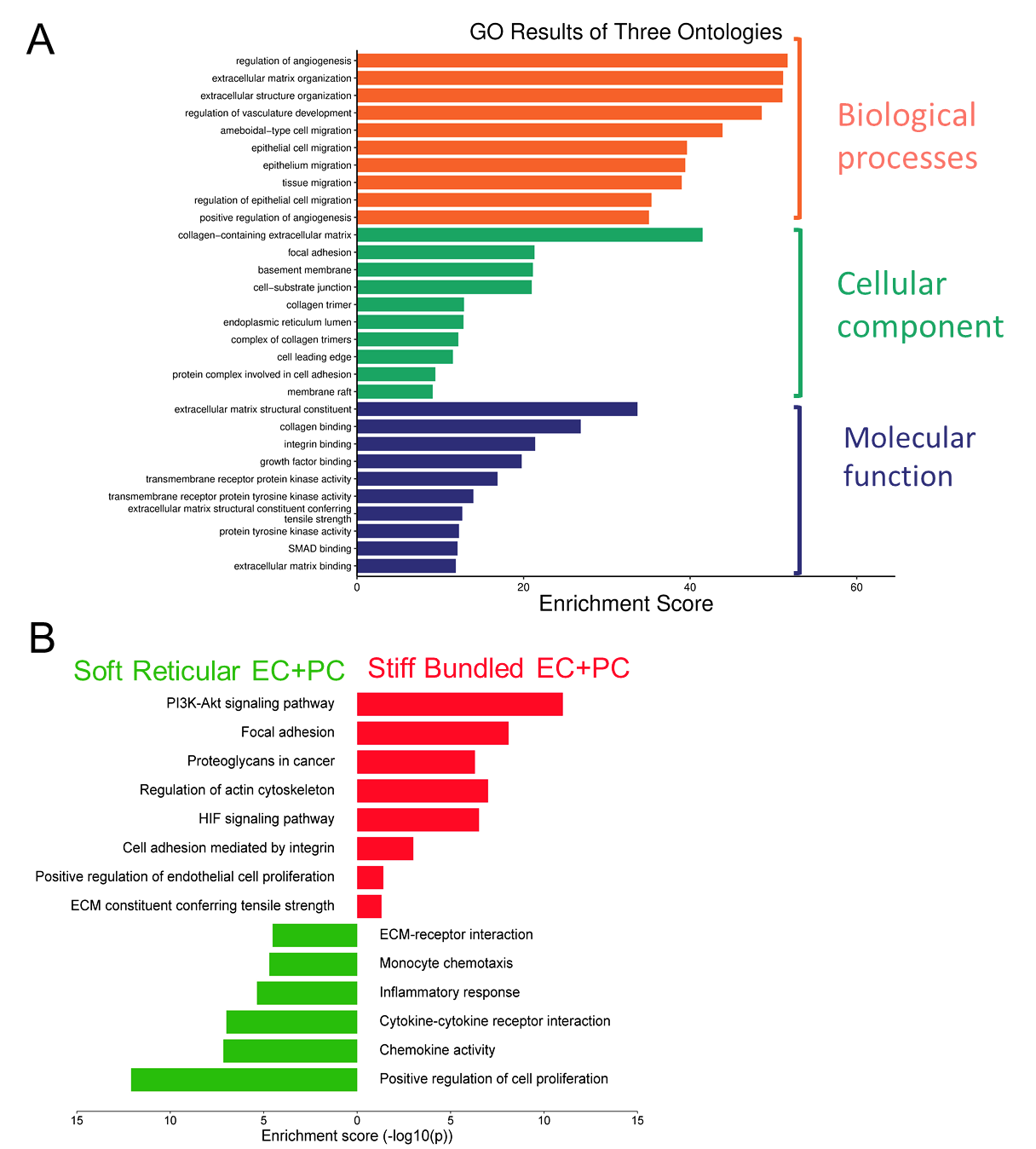

**Figure S11**. **Enrichment score of the underscored pathways comparing soft reticular and stiff bundled collagen with perivascular cells.** (A,B) Most of the pathways were related to extracellular matrix interactions and different types of cell migration, which correlate with the results observed in the immunofluorescence images.

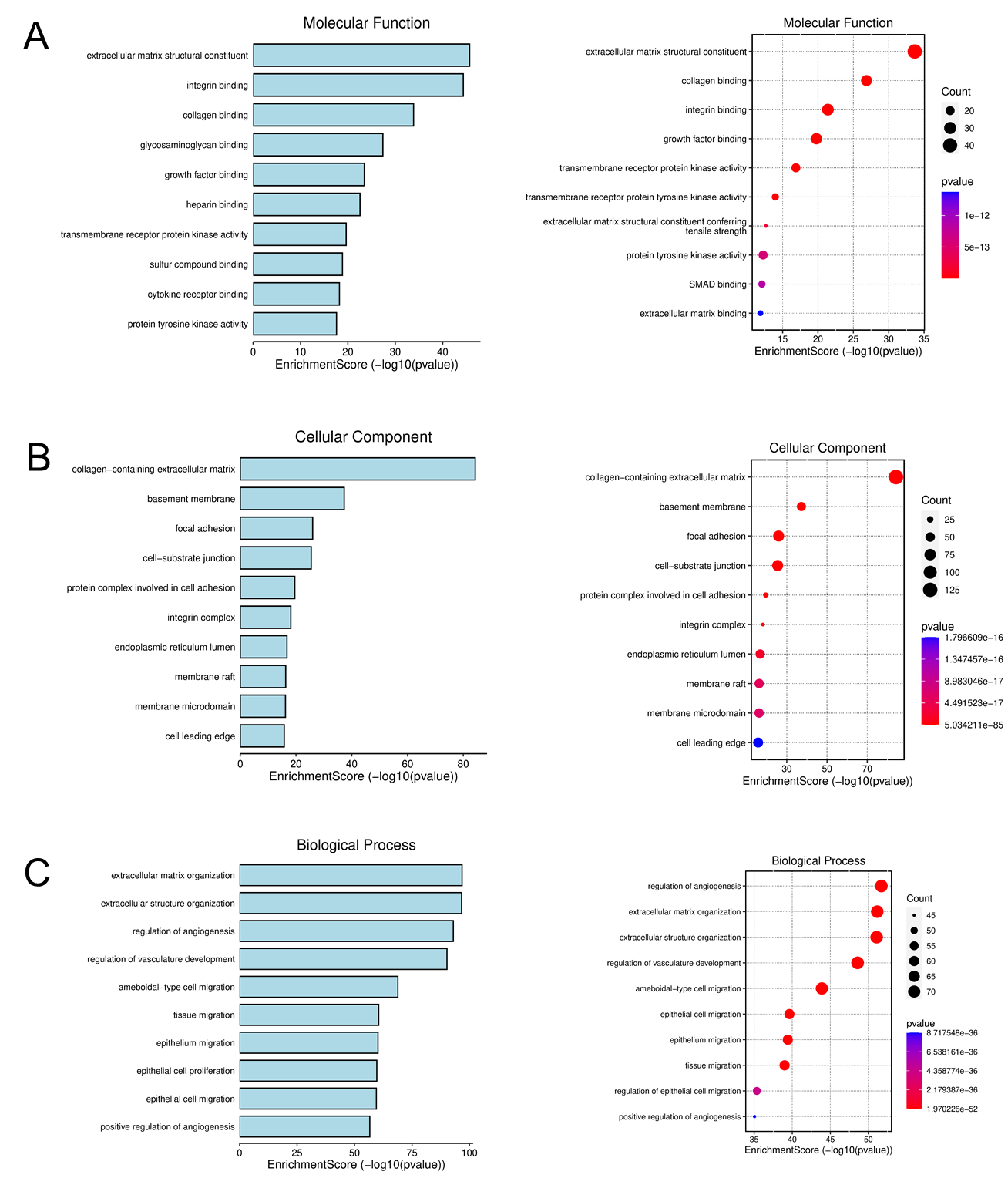

**Figure S12.Enrichment scores for capillaries engineered with perivascular cells within either soft reticular or stiff bundled collagen.** (A) molecular function, (B) cellular components and (B) biological processes show pathways related to cell-extracellular matrix binding, cell migration and angiogenesis are the most activated in the vasculature engineered in the stiff bundled collagen.

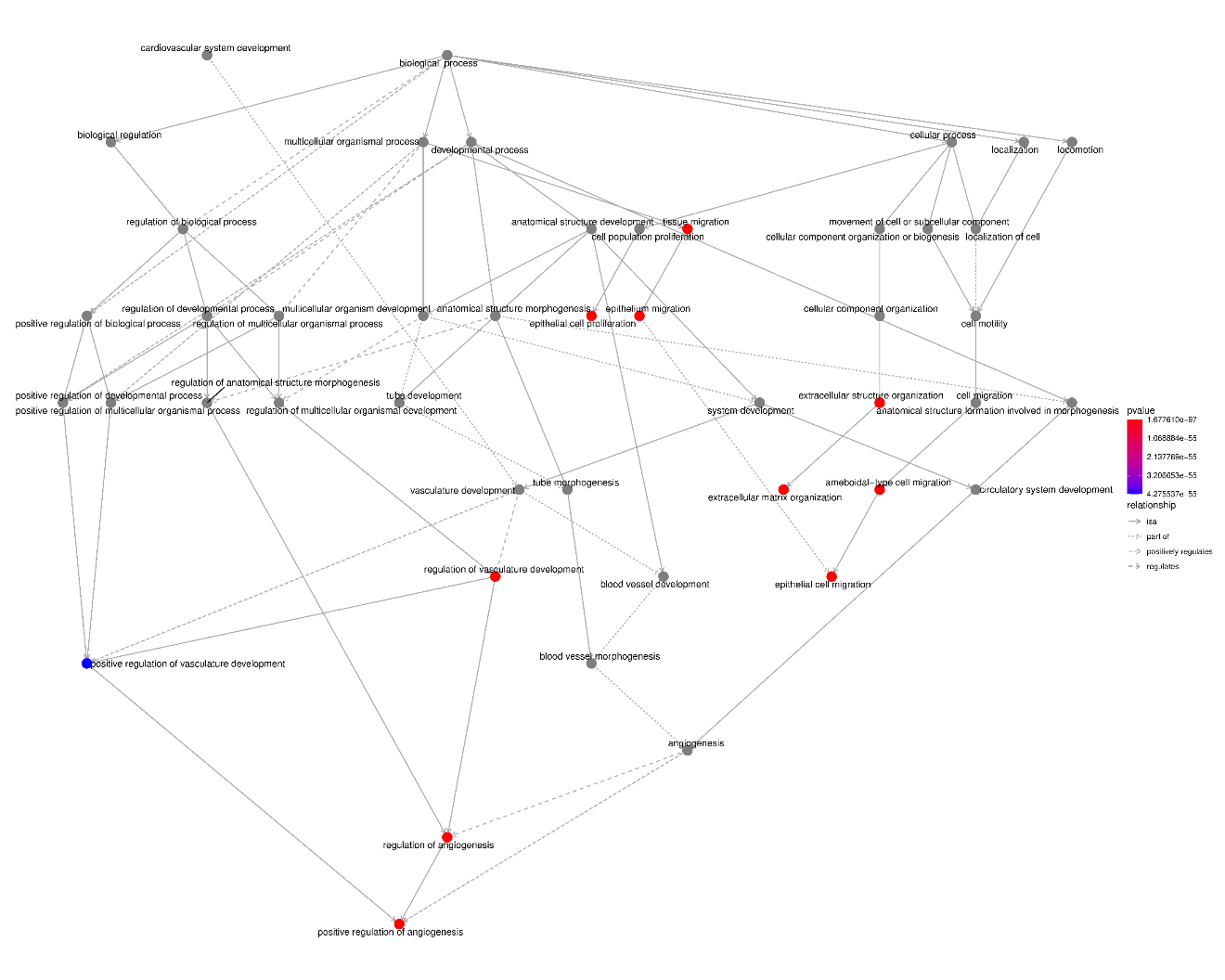

**Figure S13**. **Pathways activated in capillaries with perivascular cells within stiff bundled collagen**.

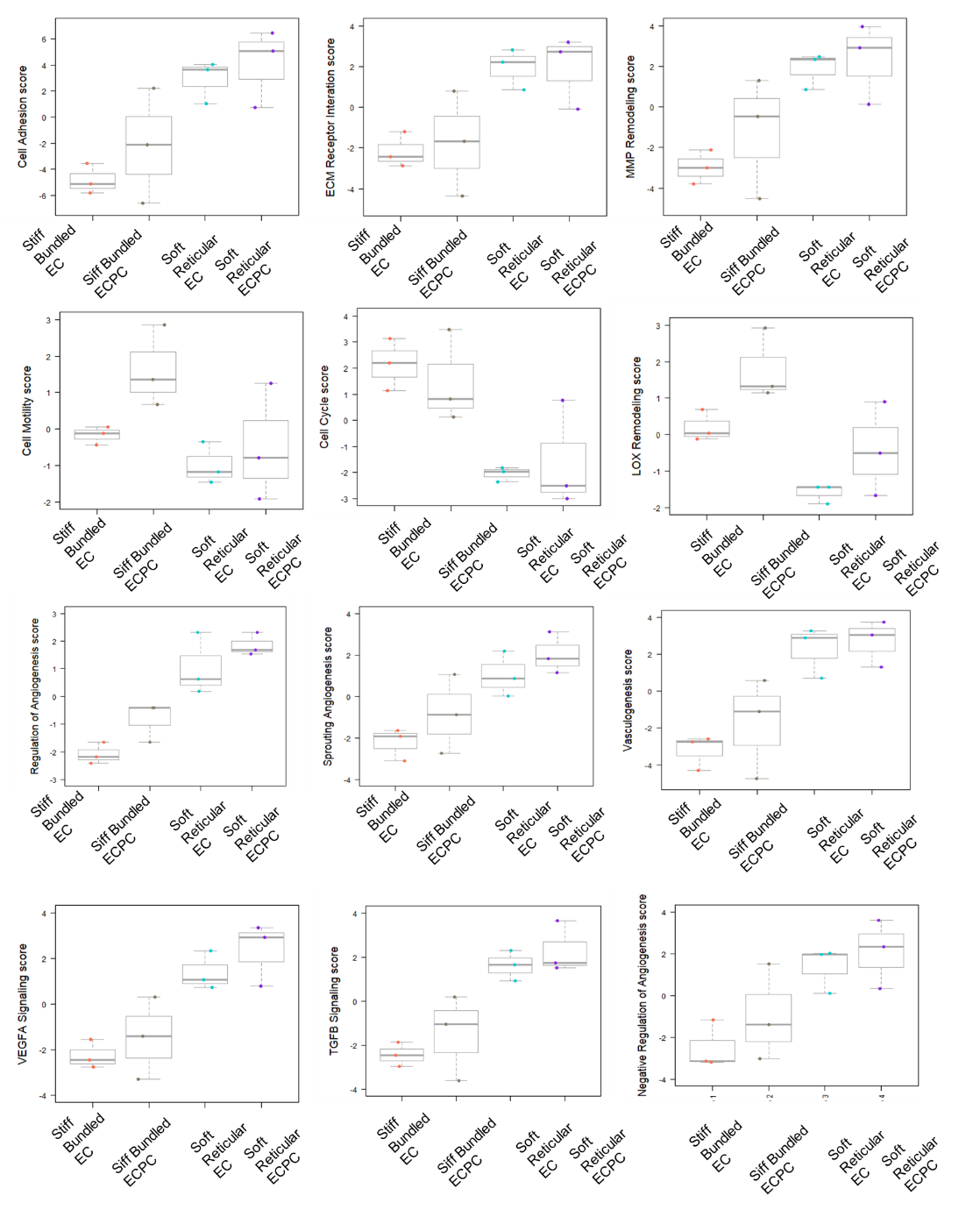

**Figure S14**. **Comparison of pathways scores among the presence of perivascular cells (PC), more fibrotic (stiff bundled) and soft reticular (healthy) collagen**. Rosalind (Nanostring ^TM^) output of data highlighting the mostly differentially expressed enrichment scores.

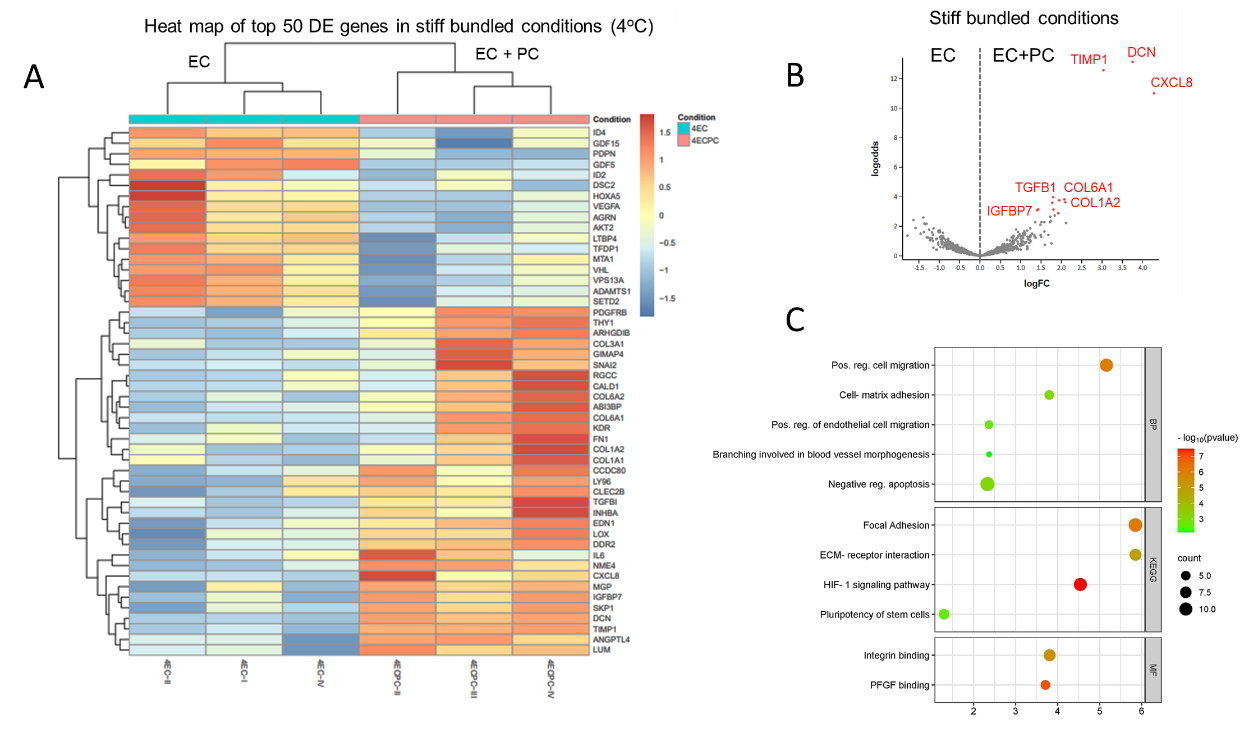

**Figure S15**. **Comparison of 50 most differentially expressed genes in stiff bundled (more fibrotic) collagen vasculature with and without perivascular cells.** (A) Heat map and (B) volcano plot highlight a prominent differential expression for *TIMP1, DCN, CXCL8, COL6A1, COL1A2, TGFB1* and *IGFBP7*. (C) The pathway analysis underscores an increase in positive regulation of cell migration, cell-matrix adhesion, focal adhesion, ECM-receptor interaction, positive regulation of endothelial cell migration, vascular branching and negative regulation of apoptosis.

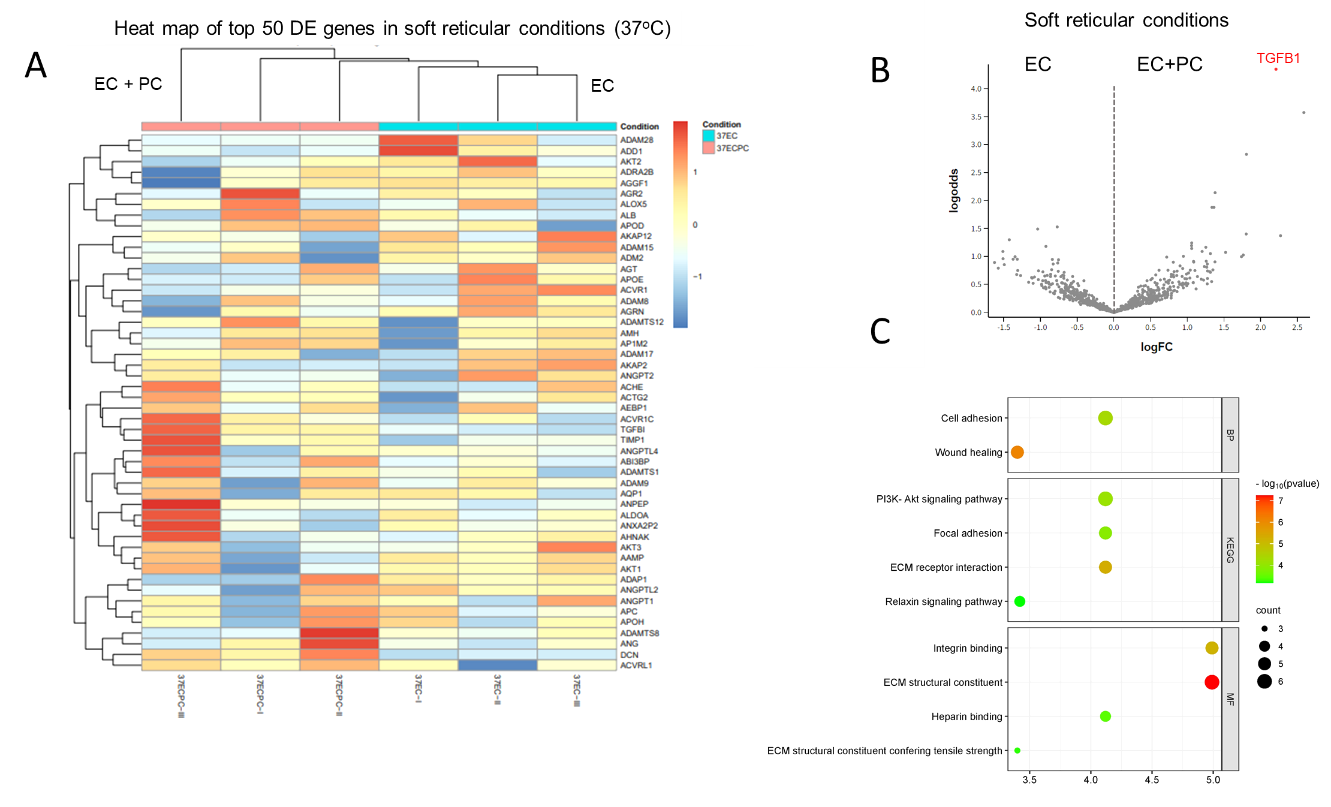

**Figure S16. Comparison of 50 most differentially expressed genes in soft reticular (healthy) collagen vasculature with and without perivascular cells.** (A) heat map and (B) volcano plot show that the mostly differentially expressed gene was transforming growth factor beta 1 (*TGFB1*). (C) The pathway analysis showed an increase in cell adhesion, focal adhesion, ECM receptor interaction, integrin binding and ECM structural component in the presence of perivascular cells.

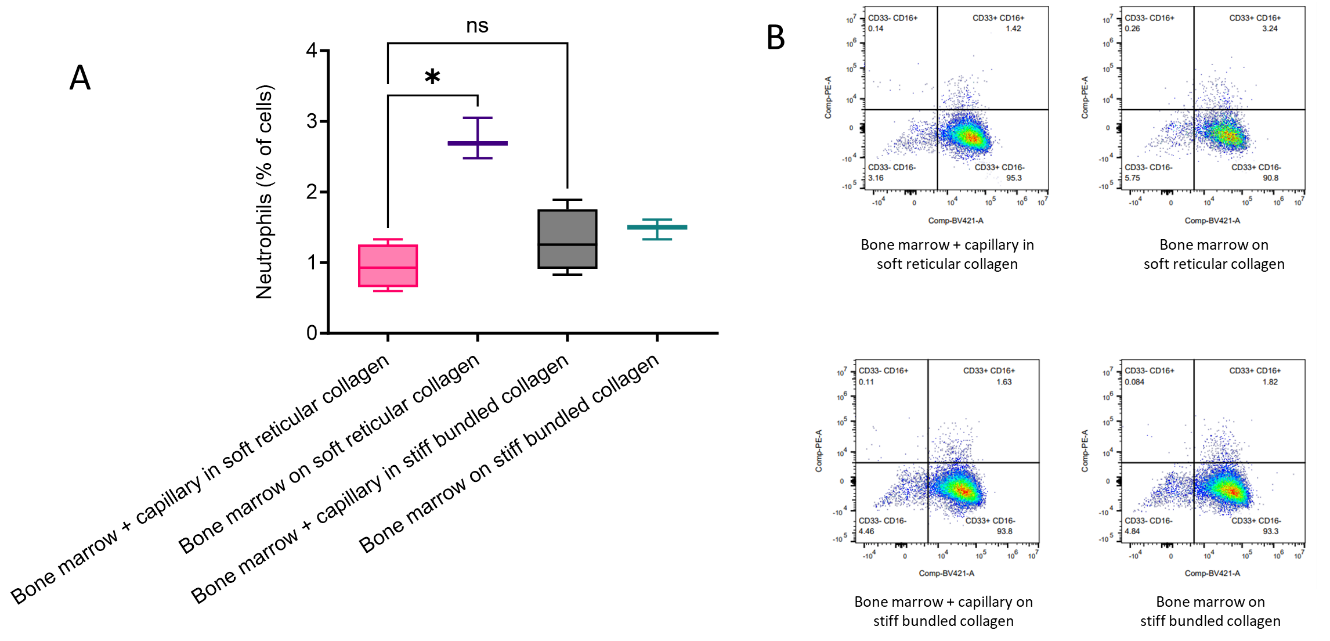

**Figure S17**. **Engineered bone marrow differentiation into neutrophilic lineage in proximity to capillaries engineered in either soft reticular or stiff bundled collagen**. (A) Flow cytometry analysis revealed comparable bone marrow cell differentiation into neutrophils regardless of collagen stiffness and microarchitecture. (B) Negative controls were performed by seeding the bone marrow on top of soft or stiff bundled collagen without the vasculature. Bone marrow seeded on top of soft reticular collagen led to an increased percentage of neutrophil differentiation.

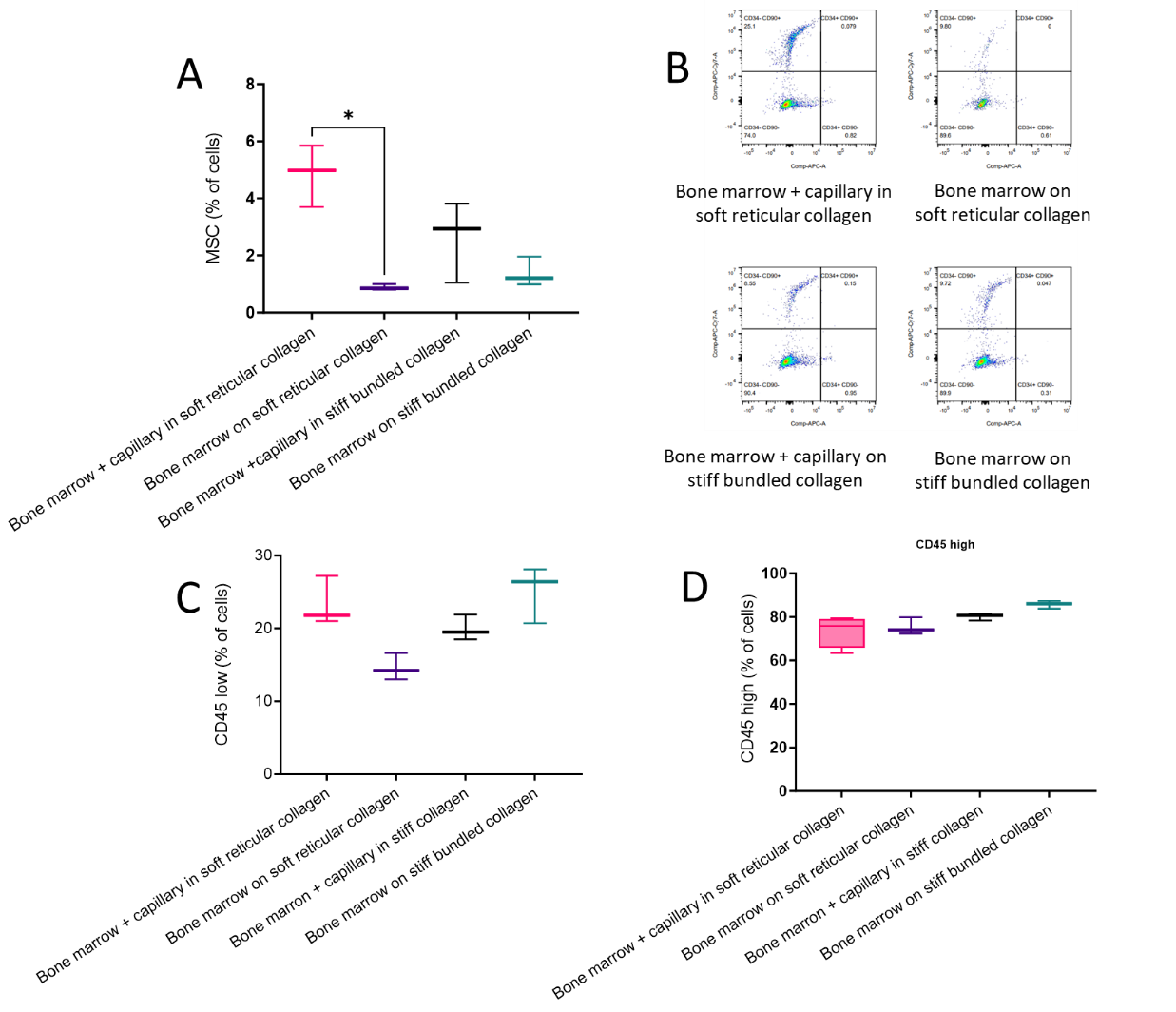

**Figure S18**. **Engineered bone marrow differentiation in proximity to healthy (soft) or more fibrotic (stiff) vasculature**. (A,B) Bone marrow seeded on top of healthy/soft collagen resulted in a decreased percentage of hMSC differentiation. (C,D) Flow cytometry analysis revealed comparable bone marrow cell differentiation toward mesenchymal stem cell lineage, as well as the presence of CD45 high or low, indicating consistent outcomes across both vascular conditions. Negative controls were performed by seeding the bone marrow on top of soft or stiff collagen without the vasculature.

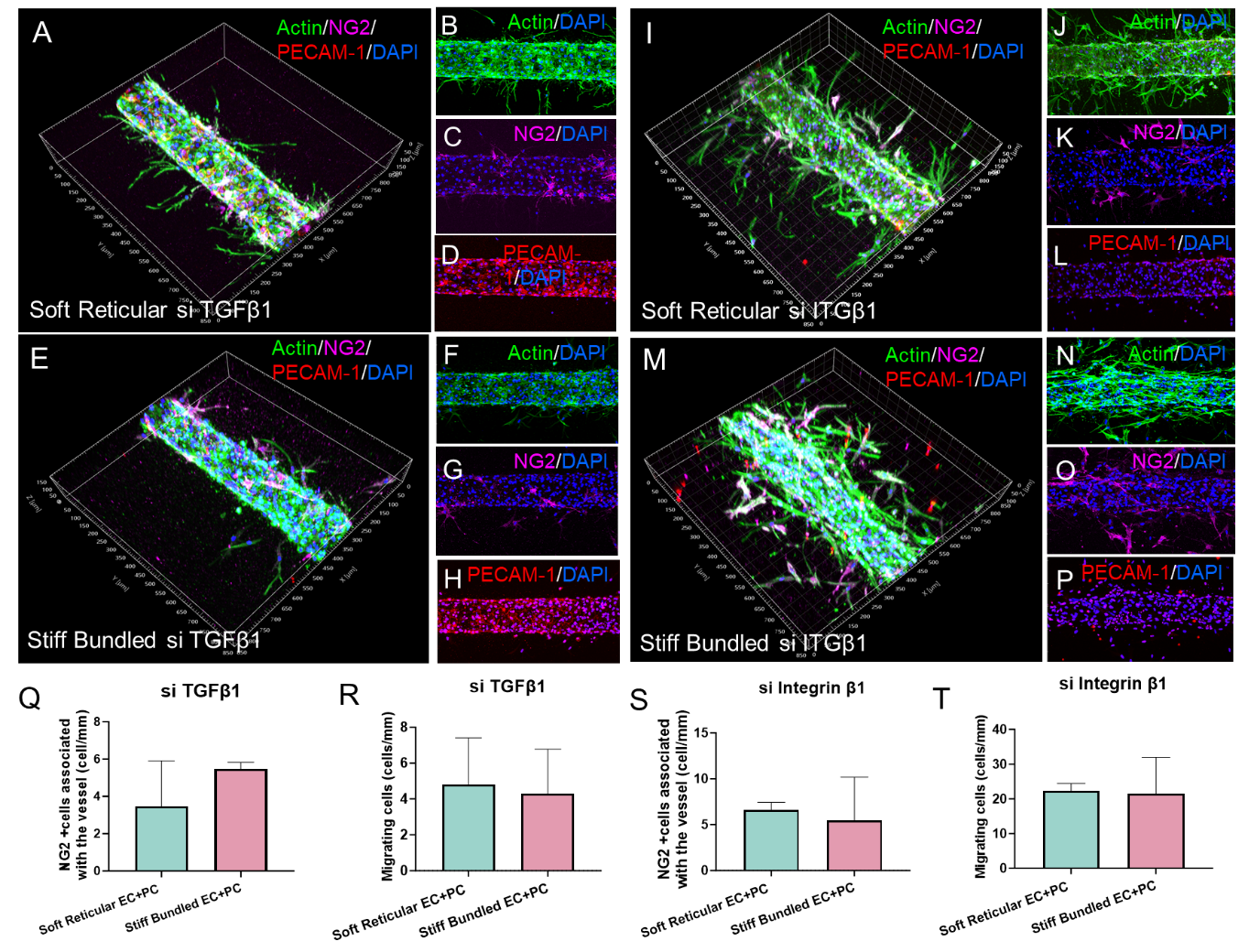

**Figure S19**. **Investigating the role of transforming growth factor 1 (TGFβ1) and integrin beta 1 (ITGβ1) in perivascular cell mechanosensing**. (A-H, Q-R) Silencing *TGFB1* gene normalizes pericyte coverage and migratory cell numbers in fibrotic vasculature, while silencing integrin B1 (*ITGB1*) (I-P, S,T) disrupts healthy vasculature, resembling fibrotic vessel traits with increased migration and reduced pericyte coverage. Note the large increase in Y-axis scaling when quantifying migrating cells in panel T.

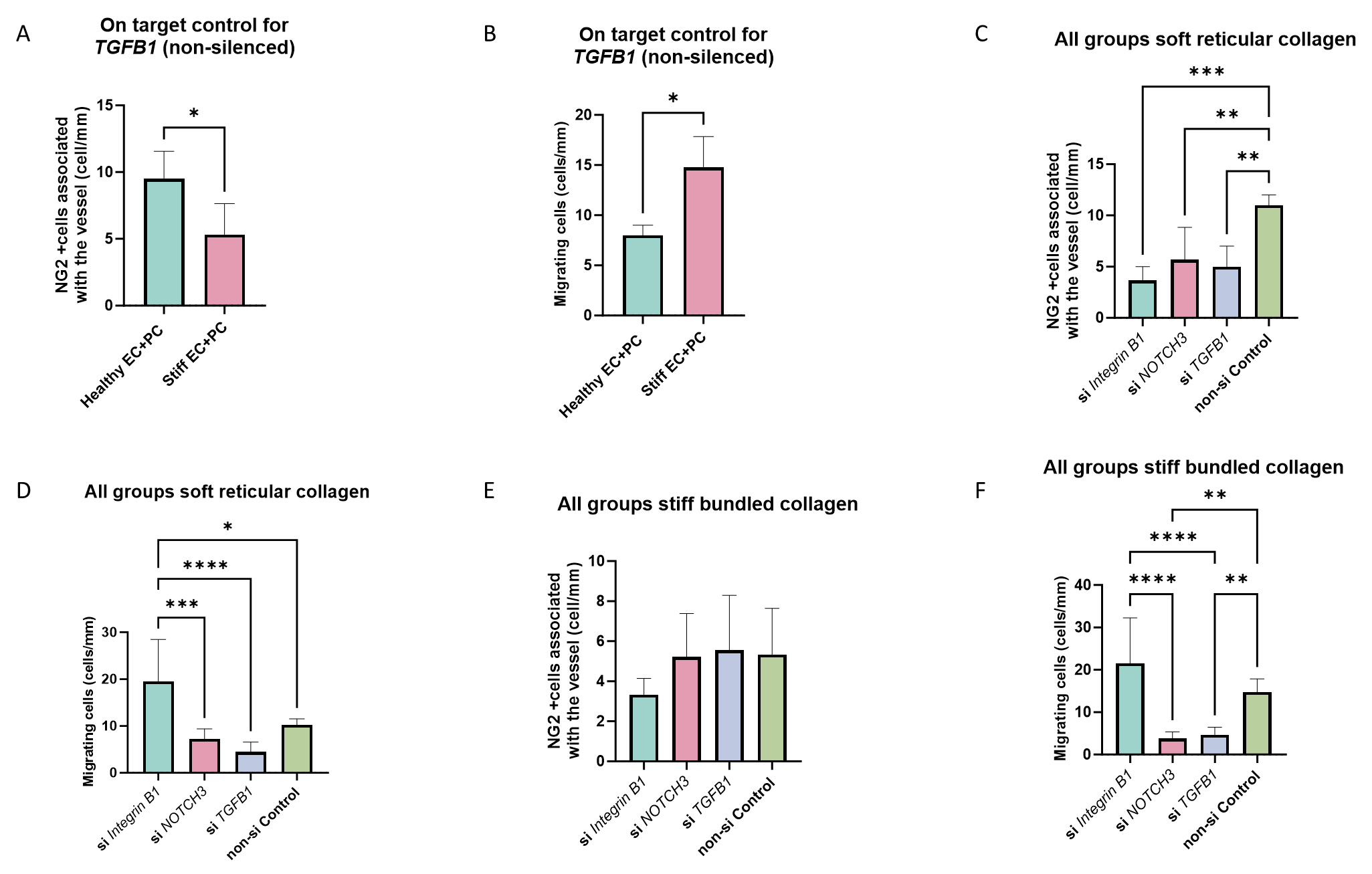

**Figure S20**. **Controls for silencing experiments.** (A,B) Non-silenced controls for *TGFB1* experiments. (C) Quantification of NG2+ cells for all silenced genes *– Integrin B1, NOTCH3* and *TGFB1* – in vasculature engineered in soft reticular collagen, showing that after silencing, all groups presented less NG2+ cells than non-silenced groups. (D) Quantification of migrating cells in the vasculature after silencing the genes demonstrating that silencing *Integrin B1* resulted in more cell migration than the control in soft reticular collagen. (E) Silencing the genes in vasculature engineered in the stiff bundled collagen did not alter the number of NG2+ cells. However, (F) migration was decreased after *NOTCH3* and *TGFB1* were silenced.

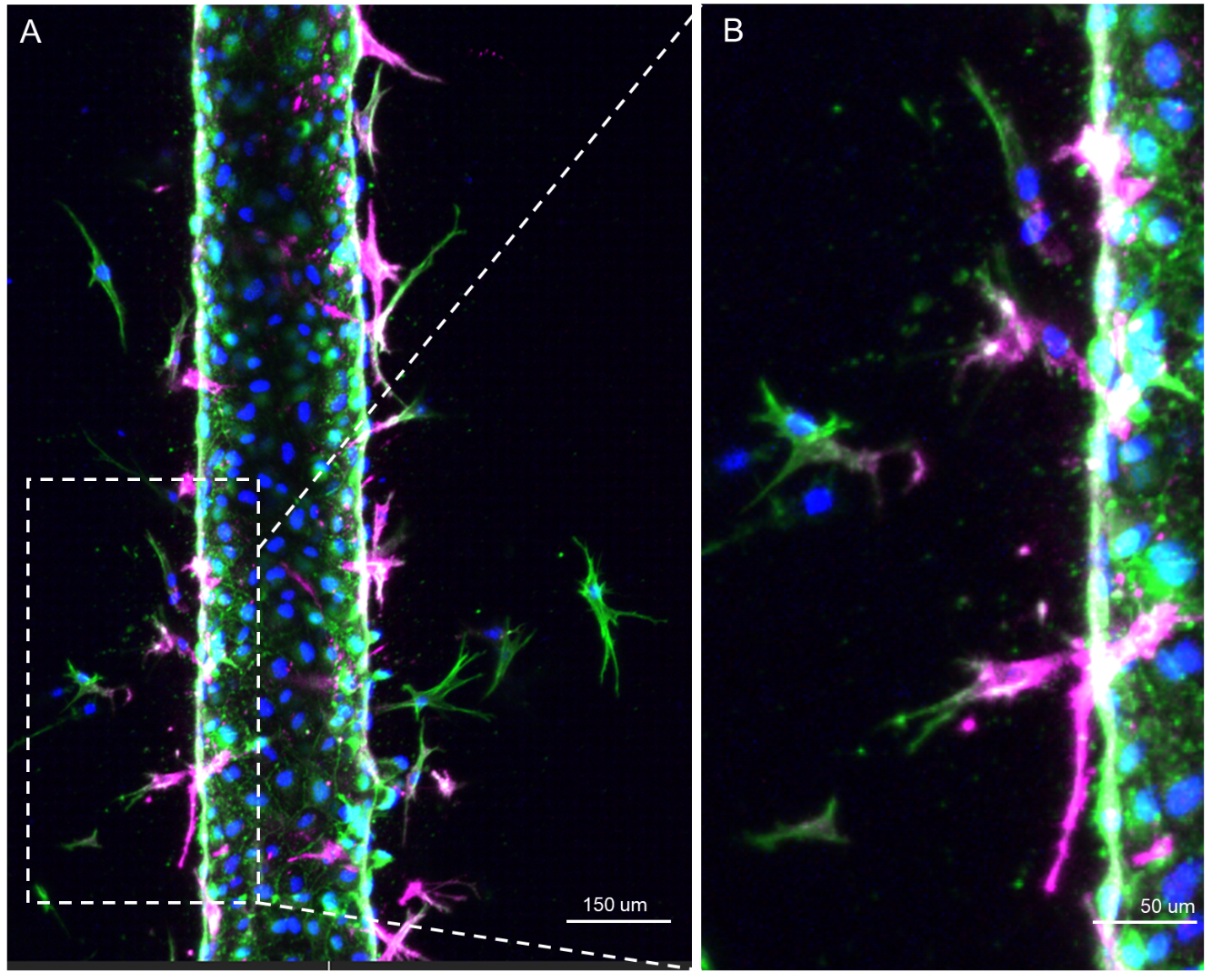

**Figure S21. Gradient of NG2 expression in perivascular cells relative to proximity to endothelial cells.** The data indicate a reduction in NG2 expression in perivascular cells that have migrated furthest from the endothelial cells, suggesting a potential juxtacrine signaling mechanism between endothelial cells and perivascular cells.

Movie S1. Capillary engineered with perivascular cells in soft reticular collagen akin healthy collagen. Cells were immunostained for actin (green), PECAM-1 (red), NG2 (magenta), DAPI (blue).

Movie S2. Capillary engineered with perivascular cells in stiff bundled collagen akin more fibrotic collagen. Cells were immunostained for actin (green), PECAM-1 (red), NG2 (magenta), DAPI (blue).

Movie S3. Barrier function assay in a healthy capillary engineered within a soft reticular collagen (37^o^C).

Movie S4. Barrier function assay in a capillary engineered within an intermediate reticular collagen (21^o^C).

Movie S5. Barrier function assay in a capillary engineered within an intermediate bundled collagen (16^o^C).

Movie S6. Barrier function assay in a capillary engineered within a fibrotic highly fibrillar collagen (4^o^C).

Movie S7. Barrier function assay in a capillary engineered within a highly reticular collagen (37^o^C) and perivascular cells with on-target control gene for *NOTCH3* (soft reticular control group).

Movie S8. Barrier function assay in a capillary engineered within a highly reticular collagen (37^o^C) and perivascular cells with silenced gene for *NOTCH3* (silenced soft reticular group).

Movie S9. Barrier function assay in a capillary engineered within a stiff bundled collagen (4^o^C) and perivascular cells with on-target control gene for *NOTCH3* (stiff bundled control group).

Movie S10. Barrier function assay in a capillary engineered within a stiff bundled (4^o^C) collagen and perivascular cells with silenced gene for *NOTCH3* (silenced stiff bundled group).
